# The user’s guide to comparative genomics with EnteroBase, including case studies on transmissions of micro-clades of *Salmonella*, the phylogeny of ancient and modern *Yersinia pestis* genomes, and the core genomic diversity of all *Escherichia*

**DOI:** 10.1101/613554

**Authors:** Zhemin Zhou, Nabil-Fareed Alikhan, Khaled Mohamed, Yulei Fan, the Agama Study Group, Mark Achtman

**Affiliations:** Warwick Medical School, University of Warwick, Gibbet Hill Road, Coventry, CV4 7AL, United Kingdom

## Abstract

EnteroBase is an integrated software environment which supports the identification of global population structures within several bacterial genera that include pathogens. Here we provide an overview on how EnteroBase works, what it can do, and its future prospects. EnteroBase has currently assembled more than 300,000 genomes from Illumina short reads from *Salmonella, Escherichia, Yersinia, Clostridiodes, Helicobacter, Vibrio*, and *Moraxella*, and genotyped those assemblies by core genome Multilocus Sequence Typing (cgMLST). Hierarchical clustering of cgMLST sequence types allows mapping, a new bacterial strain to predefined population structures at multiple levels of resolution within a few hours after uploading its short reads. Case study 1 illustrates this process for local transmissions of *Salmonella enterica* serovar Agama between neighboring social groups of badgers and humans. EnteroBase also supports SNP calls from both genomic assemblies and after extraction from metagenomic sequences, as illustrated by case study 2 which summarizes the microevolution of *Yersinia pestis* over the last 5,000 years of pandemic plague. EnteroBase can also provide a global overview of the genomic diversity within an entire genus, as illustrated by case study 3 which presents a novel, global overview of the population structure of all of the species, subspecies and clades within *Escherichia*.

## Introduction

Epidemiological transmission chains of *Salmonella, Escherichia* or *Yersinia* have been reconstructed with the help of single nucleotide polymorphisms (SNPs) from hundreds or even thousands of core genomes (Zhou *et al.* 2013; Zhou *et al.* 2014; Langridge *et al.* 2015; Connor *et al.* 2016; Dallman *et al.* 2016; Wong *et al.* 2016; Ashton *et al.* 2017; Waldram *et al.* 2017; Worley *et al.* 2018; Alikhan *et al.* 2018; Zhou *et al.* 2018c; Johnson *et al.* 2019). However, the scale of these studies pales in comparison to the numbers of publicly available archives (SRAs) of short read sequences of bacterial pathogens which have been deposited since the recent drop in price of high throughput sequencing (Wetterstrand 2019). In October 2019, the SRA at NCBI contained genomic sequence reads from 430,417 *Salmonella, Escherichia/Shigella, Clostridiodes, Vibrio* and *Yersinia*. However, until very recently (Sanaa *et al.* 2019), only relatively few draft genomic assemblies were publicly available, and even the current comparative genomic analyses in GenomeTrakr (https://www.ncbi.nlm.nih.gov/pathogens/) are restricted to relatively closely related genetic clusters. Since 2014, EnteroBase (https://enterobase.warwick.ac.uk) has attempted to address this gap for selected genera that include bacterial pathogens (Table 1). EnteroBase provides an integrated software platform (Fig. 1) that can be used by microbiologists with limited bioinformatic skills to upload short reads, assemble and genotype genomes, and immediately investigate their genomic relationships to all natural populations within those genera. These aspects have been illustrated by recent publications providing overviews of the population structures of *Salmonella* (Alikhan et al. 2018) and *Clostridioides* (Frentrup *et al*. 2019), a description of the GrapeTree GUI (Zhou *et al.* 2018a) and a reconstruction of the genomic history of the *S. enterica* Para C Lineage (Zhou *et al.* 2018c). However, EnteroBase also provides multiple additional features, which have hitherto largely been promulgated by word of mouth. Here we provide a high-level overview of the functionality of EnteroBase, followed by exemplary case studies of *Salmonella enterica* serovar Agama, *Yersinia pestis* and all of *Escherichia*.

**Table 1.**
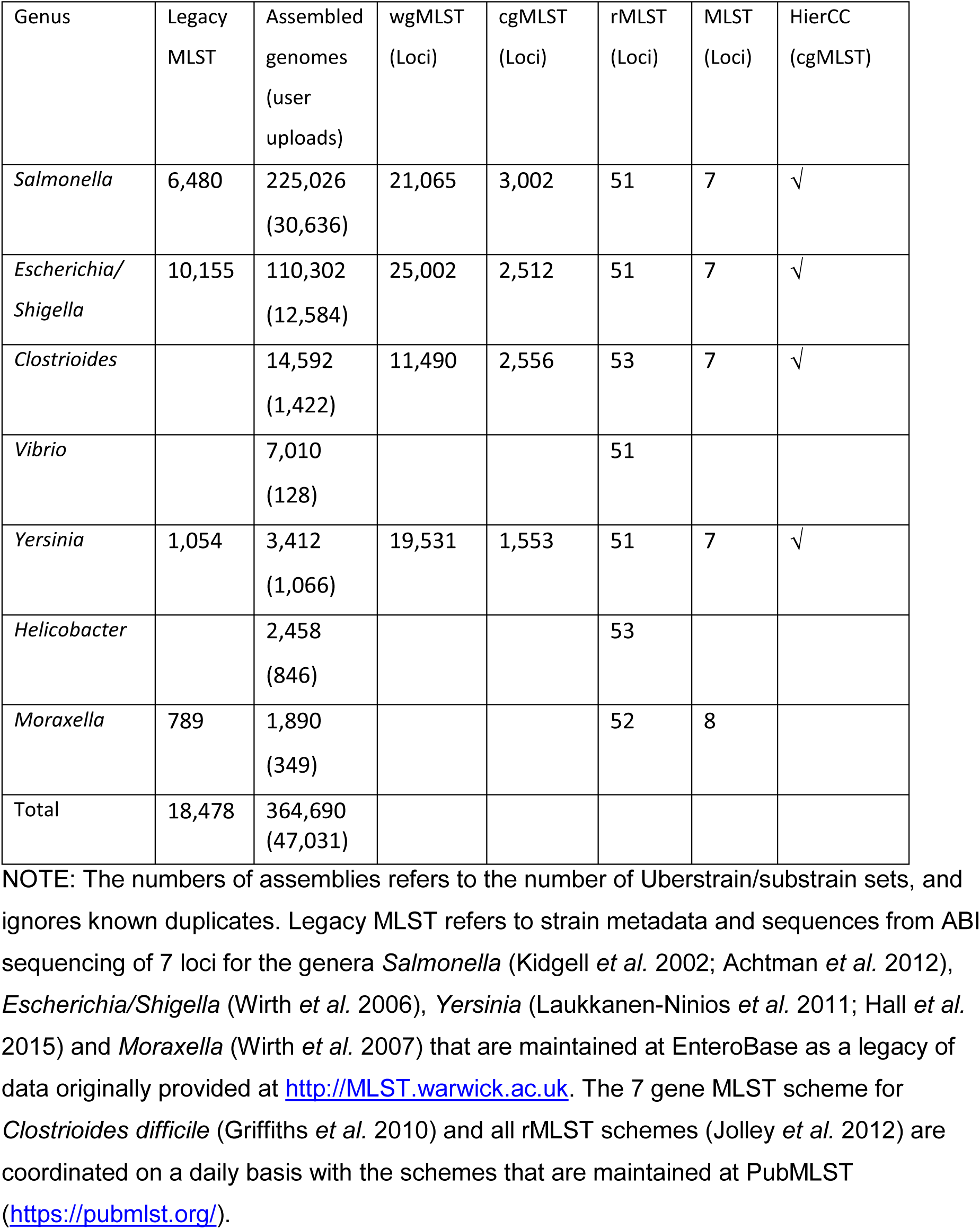
Basic statistics on EnteroBase (https://enterobase.warwick.ac.uk) (19.09.2019)

**Figure 1.**
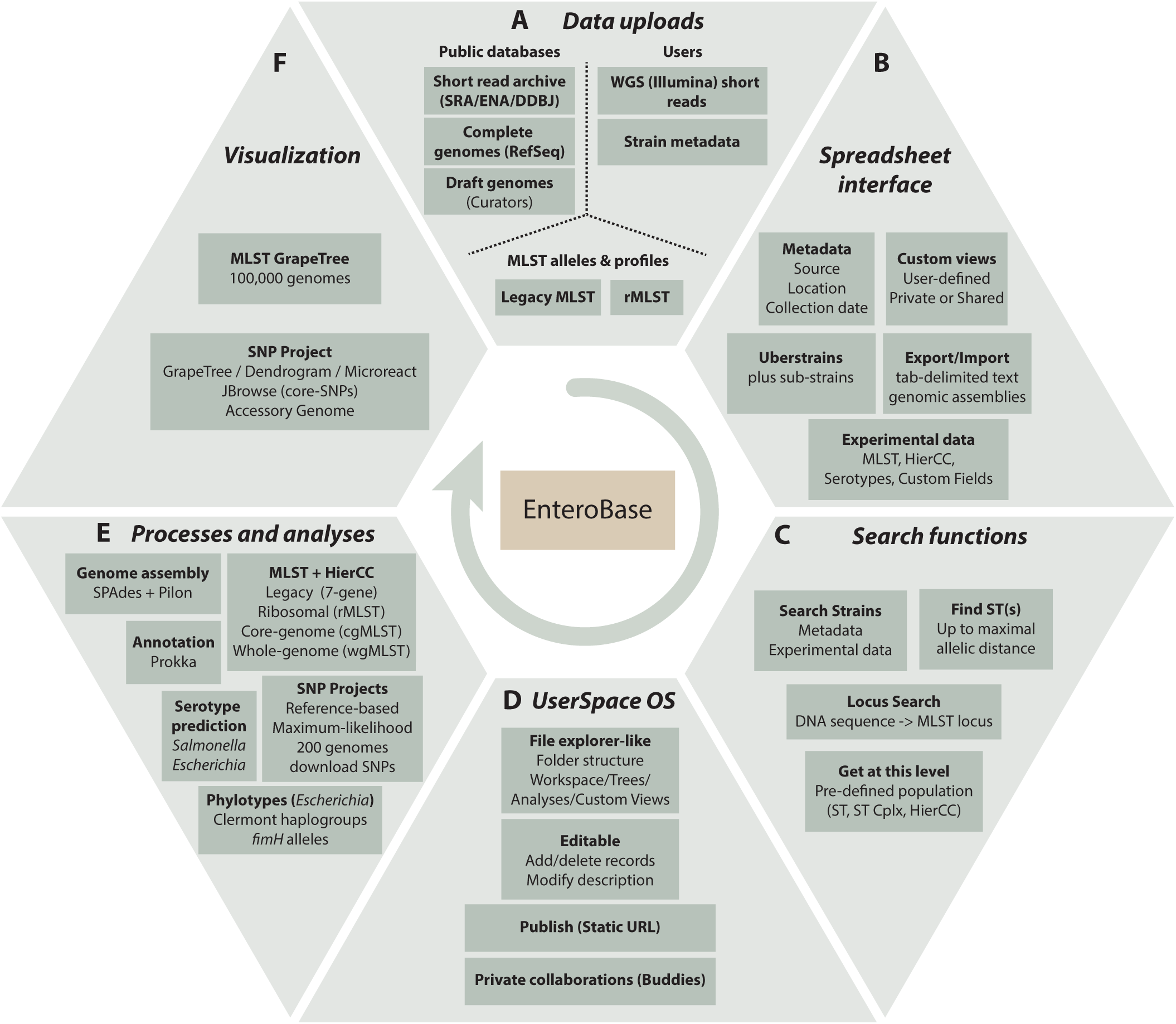
Overview of EnteroBase Features. A) Data uploads. Data are imported from public databases, user uploads and existing legacy MLST and rMLST databases at PubMLST (https://pubmlst.org/). B) Spreadsheet Interface. The browser-based interface visualizes sets of strains (one Uberstrain plus any number of sub-strains) each containing metadata, and their associated experimental data and custom views. Post-release data can be exported (downloaded) as genome assemblies or tab-delimited text files containing metadata and experimental data. Metadata can be imported to entries for which the user has editing rights by uploading tab-delimited text-files. C) Search Strains supports flexible (AND/OR) combinations of metadata and experimental data for identifying entries to load into the spreadsheet. Find ST(s) retrieves STs that differ from a given ST by no more than a maximal number of differing alleles. Locus Search uses BLASTn (Altschul et al. 1990) and UBlastP in USEARCH (Edgar 2010) to identify the MLST locus designations corresponding to an input sequence. Get at this level: menu item after right clicking on experimental MLST ST or cluster numbers. D). UserSpace OS. A file-explorer like interface for manipulations of workspaces, trees, SNP projects and custom views. These objects are initially private to their creator, but can be shared with buddies or rendered globally accessible. E) Processes and analyses, EnteroBase uses EToKi and external programs as described in Supplemental Fig. S1. F) Visualization. MLST trees are visualized with the EnteroBase tools GrapeTree (Zhou et al. 2018a) and Dendrogram, which in turn can transfer data to external websites such as MicroReact (Argimon et al. 2016).

## Results

### Overview of EnteroBase

The Enterobase back-end consists of multiple, cascading automated pipelines (Supplemental Fig. S1) which implement the multiple functions that it provides (Supplemental Fig. S2A). Many of these EnteroBase pipelines are also available within EToKi (EnteroBase ToolKit), a publicly available repository (https://github.com/zheminzhou/EToKi) of useful modules (Fig. S2B-D) that facilitate genomic assemblies, MLST typing, calling non-repetitive SNPs against a reference genome, or predicting serotypes of *E. coli* from genome assemblies (EB*Eis*).

EnteroBase performs daily scans of the GenBank SRA via its Entrez APIs (Clark et al. 2016) for novel Illumina short read sequences for each of the bacterial genera that it supports. It uploads the new reads, and assembles them (EBAssembly, Fig. S2B) into annotated draft genomes, which are published if they pass quality control (Supplemental Table S1). EnteroBase fetches the metadata associated with the records, and attempts to transcribe it automatically into Enterobase metadata format (Supplemental Tables S2, S3). During the conversion, geographic metadata are translated into structured format using the Nominatim engine offered by OpenStreetMap (OpenStreetMap contributors 2017) and the host/source metadata are assigned to pre-defined categories using a pre-trained Native Bayesian classifier implemented in the NLTK Natural Language Toolkit for Python (Bird *et al.* 2009) (Supplemental Material, Supplemental Fig. S3; estimated accuracy of 60%). Registered users can upload their own Illumina short reads and metadata into EnteroBase; these are then processed with the same pipelines.

The annotated genomes are used to call alleles for Multilocus Sequence Typing (MLST) (MLSType; Fig. S2B) and their Sequence Types (STs) are assigned to population groupings as described below. *Salmonella* serovars are predicted from the legacy MLST eBurstGroups (eBGs), which are strongly associated with individual serovars (Achtman *et al.* 2012), or by two external programs (SISTR1 (Yoshida *et al.* 2016; Robertson *et al.* 2018); SeqSero2 (Zhang *et al.* 2019)) which evaluate genomic sequences. *Escherichia* serotypes are predicted from the genome assemblies by the EnteroBase module EB*Eis.* Clermont haplogroups are predicted for *Escherichia* by two external programs (ClermonTyping (Beghain *et al.* 2018); EZClermont (Waters *et al.* 2018)) and *fimH* type by a third (FimTyper (Roer *et al.* 2017)). By default, public free access to strain metadata and the genome assemblies, predicted genotypes and predicted phenotypes is immediate, but a delay in the release date of up to 12 months can be imposed when uploading short read sequences.

In September 2019, EnteroBase provided access to 364,690 genomes and their associated metadata and predictions (Table 1). In order to allow comparisons with historical data, EnteroBase also maintains additional legacy 7-gene MLST assignments (and metadata) that were obtained by classical Sanger sequencing from 18,478 strains,

#### Ownership, permanence, access and privacy

EnteroBase users can upload new entries, consisting of paired-end Illumina short reads plus their metadata. Short reads are deleted after genome assembly, or after automated, brokered uploading of the reads and metadata to the European Nucleotide Archive upon user request.

The search and graphical tools within EnteroBase include all assembled genomes and their metadata, even if they are pre-release. However, ownership of uploaded data remains with the user, and extends to all calculations performed by EnteroBase. Only owners and their buddies, admins or curators can edit the metadata. And only those individuals can download any data or calculations prior to their release date. In order to facilitate downloading of post-release data by the general community, downloads containing metadata and genotypes or genomic assemblies are automatically stripped of pre-release data for users who lack ownership privileges. Similarly, pre-release nodes within trees in the GrapeTree and Dendrogram graphical modules must be hidden before users without ownership privileges can download those trees.

In general, metadata that were imported from an SRA are not editable, except by admins and curators. However, the admins can assign editing rights to users with claims to ownership or who possess special insights.

#### MLST Population structures

Each unique sequence variant of a gene in an MLST scheme is assigned a unique numerical designation. 7-gene MLST STs consist of seven integers for the alleles of seven housekeeping gene fragments (Maiden *et al.* 1998). rSTs consist of 51-53 integers for ribosomal protein gene alleles (Jolley *et al.* 2012). cgMLST STs consist of 1,553 – 3,002 integers for the number of genes in the soft core genome for that genus (Table 1), which were chosen as described elsewhere (Frentrup *et al.* 2019). However, STs are arbitrary constructs, and natural populations can each encompass multiple, related ST variants. Therefore, 7-gene STs are grouped into ST Complexes in *Escherichia*/*Shigella* (Wirth et al. 2006) by an eBurst approach (Feil *et al.* 2004), and into their equivalent eBurst groups (eBGs) in *Salmonella* (Achtman et al. 2012). EnteroBase implements similar population groups (reBGs) for rMLST in *Salmonella*, which are largely consistent with eBGs or their sub-populations (Alikhan *et al.* 2018). The EnteroBase Nomenclature Server (Fig. S1) calculates these population assignments automatically for each novel ST on the basis of single linkage clustering chains with maximal pairwise differences of one allele for 7-gene MLST and two alleles for rMLST. In order to prevent overlaps between ST Complexes, growing chains are terminated when they extend too closely to other existing populations (2 alleles difference in 7-gene MLST and 5 in rMLST).

cgMLST has introduced additional complexities over MLST and rMLST. Visual comparisons of cgSTs are tedious, and rarely productive, because each consists of up to 3,002 integers. Furthermore, almost all cgSTs contain some missing data because they are called from draft genomes consisting of multiple contigs. EnteroBase contains 100,000s of cgST numbers because almost every genome results in a unique cgST number, even though many cgSTs only differ from others by missing data. EnteroBase supports working with so many cgSTs through HierCC (Hierarchical Clustering), a novel approach which supports analyses of population structures based on cgMLST at multiple levels of resolution. In order to identify the cut-off values in stepwise cgMLST allelic distances which would reliably resolve natural populations, we first calculated a matrix of pair-wise allelic distances (excluding pairwise missing data) for all existing pairs of cgSTs, and one matrix for the HierCC clusters at each level of allelic distance, i.e. one matrix for HC0, HC1, HC2…HC3,001. A genus-specific subset of the most reliable HierCC clusters is reported by EnteroBase.

For *Salmonella*, thirteen HierCC levels are reported, ranging from HC0 (indistinguishable except for missing data) to HC2850 (Fig. 2). Our experience with *Salmonella* indicates that HC2850 corresponds to subspecies, HC2000 to super-lineages (Zhou *et al.* 2018c) and HC900 to cgMLST versions of eBGs. Long-term endemic persistence seems to be associated with HC100 or HC200; and epidemic outbreaks with HC2, HC5 or HC10. Eleven levels are reported for the other genera, ranging from HC0 up to HC2350 for *Escherichia*, HC2500 for *Clostridioides* and HC1450 for *Yersinia. Escherichia* HC1100 corresponds to ST Complexes (man. In prep.) and the correspondences to population groupings in *Clostridioides* are described elsewhere (Frentrup et al. 2019). Further information on HierCC can be found in the EnteroBase documentation (https://tinyurl.com/HierCC-doc).

**Figure 2.**
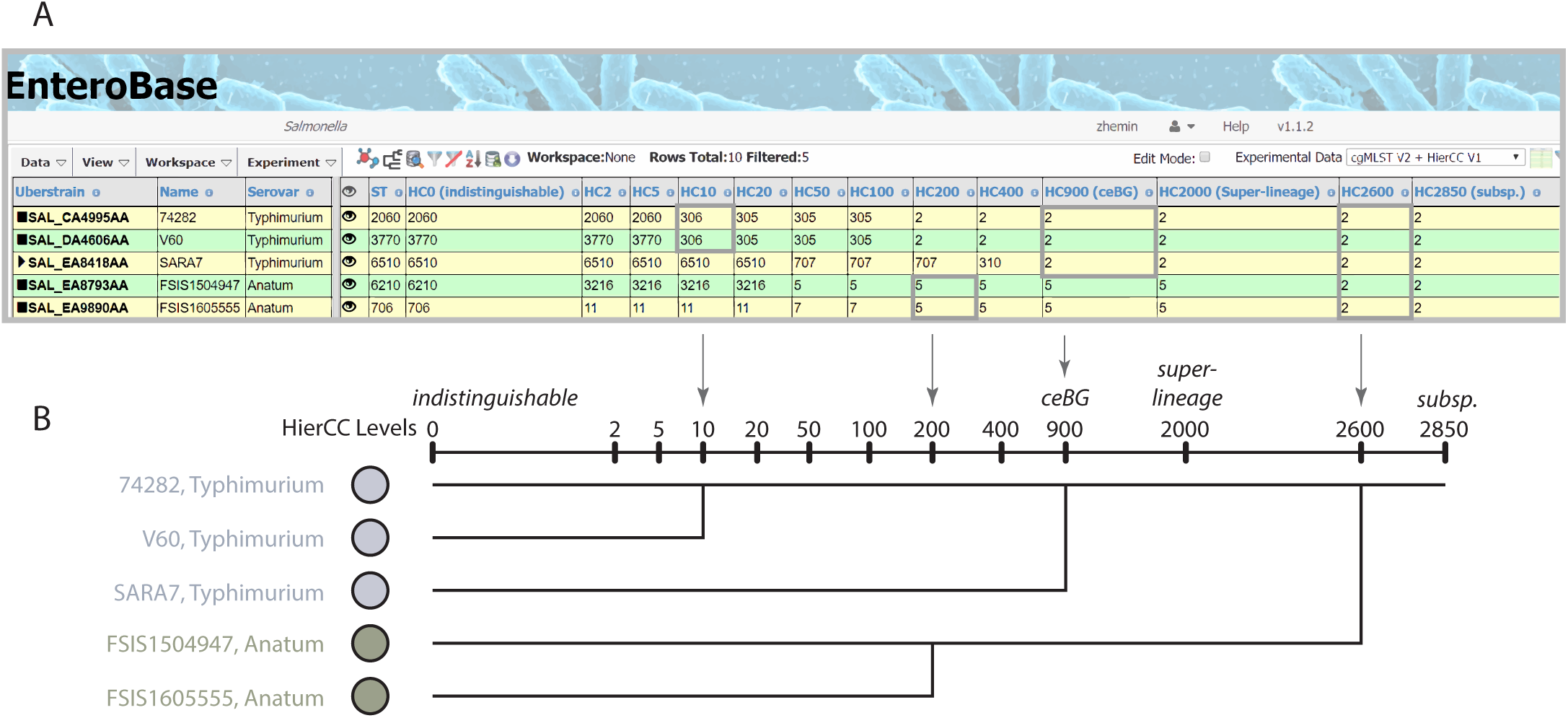
The hierarchical cgMLST clustering (HierCC) scheme in Enterobase. A) A screenshot of *Salmonella* cgMLST V2 plus HierCC V1 data for five randomly selected genomes. The numbers in the columns are the HierCC cluster numbers. Cluster numbers are the smallest cgMLST ST number in single-linkage clusters of pairs of STs that are joined by up to the specified maximum number of allelic differences. These maximum differences are indicated by the suffix of each HC column, starting with HC0 for 0 cgMLST allelic differences through to HC2850 for 2850 allelic differences. The cluster assignments are greedy because individual nodes which are equidistant from multiple clusters are assigned to the cluster with the smallest cluster number. B) Interpretation of HierCC numbers. The assignments of genomic cgMLST STs to HC levels can be used to assess their genomic relatedness. HC0 indicates identity except for missing data. The top two genomes are both assigned to HC10_306, which indicates a very close relationship, and may represent a transmission chain. The top three genomes are all assigned to HC900_2, which corresponds to a legacy MLST eBG. HC2000 marks super-lineages (Zhou *et al*. 2018c) and HC2850 marks subspecies. This figure illustrates these interpretations in the form of a cladogram drawn by hand.

#### Uber- and Sub-strains

Most bacterial isolates/strains in EnteroBase are linked to one set of metadata and one set of genotyping data. However, EnteroBase includes strains for which legacy MLST data from classical Sanger sequencing exists in addition to MLST genotypes from genomic assemblies. Similarly, some users have uploaded the same reads to both EnteroBase and SRAs, and both sets of data are present in EnteroBase. In other cases, genomes of the same strain have been sequenced by independent laboratories, or multiple laboratory variants have been sequenced that are essentially indistinguishable (e.g. *S. enterica* LT2 or *E. coli* K-12).

EnteroBase deals with such duplications by implementing the concept of an Uberstrain, which can be a parent to one or more identical sub-strains. Sub-strains remain invisible unless they are specified in the search dialog (Supplemental Fig. S4), in which case they are shown with a triangle in the Uberstrain column (Fig. 3A). Examples of the usage of this approach can be found in Supplemental Material.

**Figure 3.**
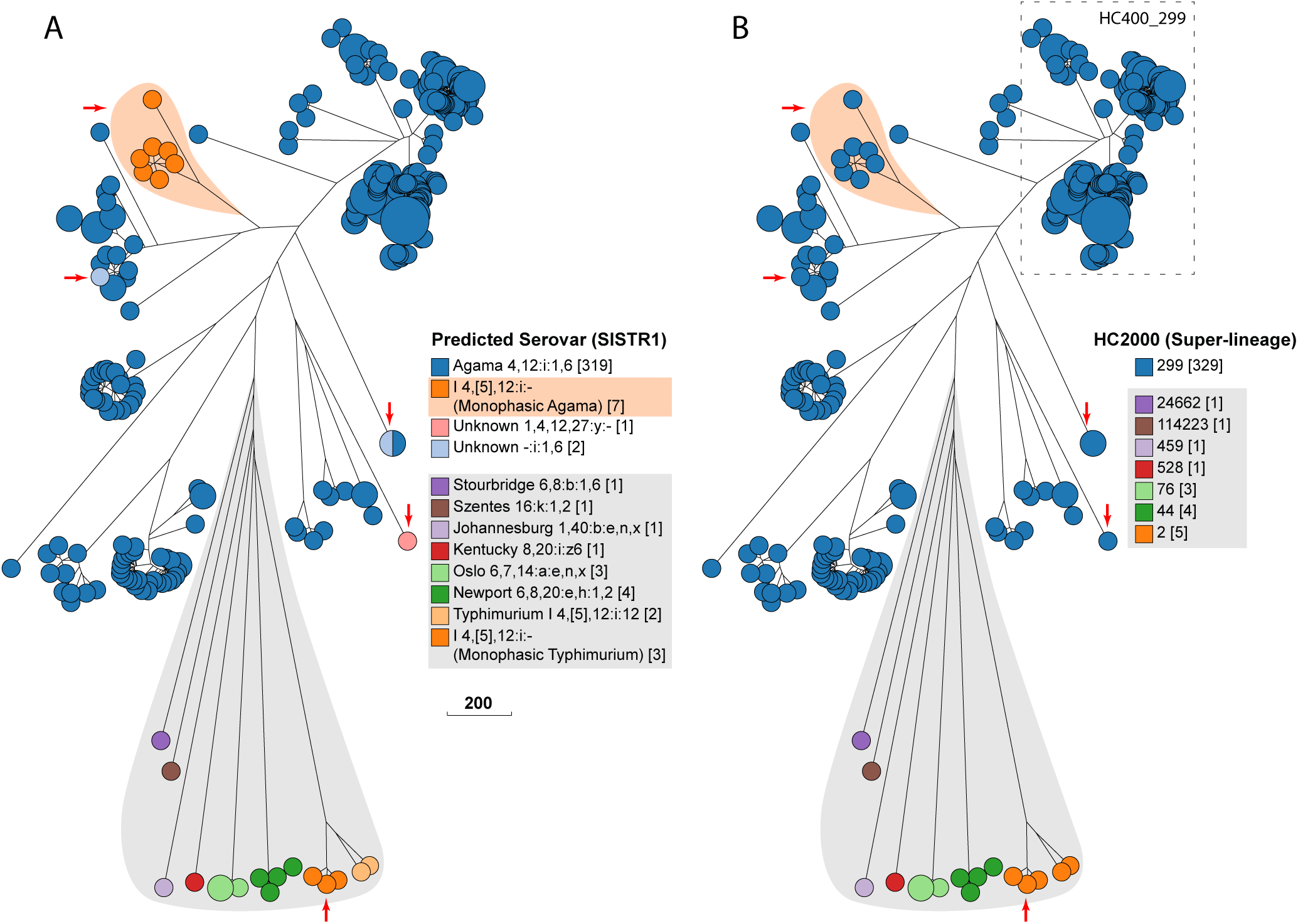
Serovar *versus* HierCC clustering in serovar Agama. GrapeTree (Zhou et al. 2018a) depiction of a Rapid-NJ tree (Simonsen *et al*. 2011) of cgMLST allelic distances between genomic entries whose metadata Serovar field contained Agama or SISTR1 (Robertson *et al*. 2018) Serovar predictions contained Agama. A) Color coding by Predicted Serovar (SISTR1). Arrows indicate isolates whose serovar was not predicted. Orange shading emphasizes 1,4,[5],12:i:- isolates that were monophasic Agama. Gray shading indicates isolates with incorrect Serovar metadata, including 1,4,[5],12:i:- isolates that were monophasic Typhimurium (arrow). B) Color-coding by HC2000 cluster. All Agama entries are HC2000_299, as were the genetically related entries marked with arrows or emphasized by orange shading. Entries from other serovars (gray shading) were in diverse other HC2000 clusters. The dashed box indicates a subset of Agama strains within HC400_299, including all isolates from badgers, which were chosen for deeper analyses in Fig. 4. Scale bar: number of cgMLST allelic differences.

### Examples of the utility of EnteroBase

Often the utility of a tool first becomes clear through examples of its use. Here we present three case studies that exemplify different aspects of EnteroBase. Case study 1 demonstrates how geographically separated laboratories can collaborate in private on an EnteroBase project until its completion, upon which EnteroBase publishes the results. This example focuses on geographical micro-variation and transmission chains between various host species of a rare serovar of *S. enterica*. Case study 2 demonstrates how to combine modern genomes of *Yersinia pestis* with partially reconstructed genomes from ancient skeletons of plague victims. It also demonstrates how EToKI can extract SNPs from metagenomic sequence reads. Case study 3 provides a quantitative overview of the genomic diversity of an entire genus, thereby defining the EcoRPlus set of representative genomes of all *Escherichia*.

### Case Study 1: A group collaboration on *S. enterica* serovar Agama

*S. enterica* subsp. *enterica* encompasses more than 1,586 defined serovars (Guibourdenche *et al.* 2010; Issenhuth-Jeanjean *et al.* 2014). These differ in the antigenic formulas of their lipopolysaccharide (O antigen) and/or two alternative flagellar antigens (H1, H2), which are abbreviated as O:H1:H2. Some serovars are commonly isolated from infections and the environment, and have been extensively studied. Others are rare, poorly understood and often polyphyletic (Achtman *et al.* 2012), including *Salmonella* that colonize badgers (Wray *et al.* 1977; Wilson *et al.* 2003).

In late 2018, serovar Agama (antigenic formula: 4,12:i:1,6) was specified in the Serovar metadata field for only 134/156,347 (0.09%) genome assemblies in EnteroBase, and all 134 isolates were from humans. We were therefore interested to learn that the University of Liverpool possessed serovar Agama isolates that had been isolated in 2006-2007 from European badgers (*Meles meles*) in Woodchester Park, Gloucestershire, England. We sequenced the genomes of 72 such isolates, and uploaded the short reads and strain metadata into EnteroBase. This data was used to analyze the population structure of a rare serovar within a single host species over a limited geographical area, and to compare Agama genomes from multiple hosts and geographical sources.

#### Search Strains

The browser interface to EnteroBase is implemented as a spreadsheet-like window called a “Workspace” that can page through 1,000s of entries, showing metadata at the left and experimental data at the right (https://tinyurl.com/EnteroBase-WS). However visual scanning of 1,000s of entries is inefficient. EnteroBase therefore offers powerful search functions (https://tinyurl.com/EnteroBase-search) for identifying isolates that share common phenotypes (metadata) and/or genotypes (experimental data).

EnteroBase also predicts serovars from assembled *Salmonella* genomes and from MLST data. However, the software predictions are not fail-proof, and many entries lack metadata information, or the metadata is erroneous. We therefore used the Search Strains dialog box to find entries containing “Agama” in the metadata field or by the predictions from SISTR1. Phylogenetic analyses of the cgMLST data from those entries indicated that Agama consisted of multiple micro-clusters.

#### International participation in a collaborative network

Almost all Agama isolates in EnteroBase were from England, which represents a highly skewed geographical sampling bias that might lead to phylogenetic distortions. We therefore formed the Agama Study Group, consisting of colleagues at national microbiological reference laboratories in England, Scotland, Ireland, France, Germany and Austria. The participants were declared as ‘buddies’ within EnteroBase (https://tinyurl.com/EnteroBase-buddies) with explicit rights to access the Workspaces and phylogenetic trees in the Workspace\Load\Shared\Zhemin\Agama folder. After completion of this manuscript, that folder was made publicly available.

We facilitated the analysis of the Agama data by creating a new user-defined Custom View (https://tinyurl.com/EnteroBase-customview), which can aggregate various sources of experimental data as well as User-defined Fields. The Custom View was saved in the Agama folder, and thereby shared with the Study Group. It too was initially private but became public together with the other workspaces and trees when the folder was made public.

Members of the Agama Study Group were requested to sequence genomes from all Agama strains in their collections, and to upload those short reads to EnteroBase, or to send their DNAs to University of Warwick for sequencing and uploading. The new entries were added to the ‘All Agama Strains’ workspace. The final set of 345 isolates had been isolated in Europe, Africa and Australia, with collection years ranging from 1956 to 2018 (Supplemental Table S3).

#### Global population Structure of Agama

We created a neighbor joining GrapeTree (Zhou *et al.* 2018a) of cgMLST data to reveal the genetic relationships within serovar Agama. Color coding the nodes of the tree by SISTR1 serovar predictions confirmed that most isolates were Agama (Fig. 3A). However, one micro-cluster (shaded in light orange) consisted of seven monophasic Agama isolates with a defective or partial *fljB* (H2) CDS, which prevented a serovar prediction. SISTR1 also could not predict the O antigens of three other related isolates (arrows in Fig. 3). Sixteen other isolates on long branches were assigned to other serovars by SISTR1 (Fig. 3A, grey shading). Comparable results were obtained with SeqSero2 or eBG serovar associations, and these sixteen isolates represent erroneous Serovar assignments within the metadata. Interestingly, three of these erroneous Agama had the same predicted antigenic formula (1,4,[5],12:i:-) as the monophasic Agama isolates (orange shading), but these represent monophasic Typhimurium.

In contrast to serovar, coloring the tree nodes by HC2000 clusters (Fig. 3B) immediately revealed that all genomes that were called Agama by SISTR1 belonged to HC2000 cluster number 299 (HC2000_299), and all HC2000_299 were genetically related and clustered together in the tree (Fig. 3B). In contrast, the 16 other isolates on long branches (gray shading) belonged to other HC2000 clusters.

These results show that Agama belongs to one super-lineage, HC2000_299, which has been isolated globally from humans, badgers, companion animals and the environment since at least 1956. The genetic relationships would not have been obvious with lower resolution MLST: some Agama isolates belong to eBG167, others to eBG336 and thirteen Agama MLST STs do not belong to any eBG.

#### Transmission patterns at different levels of HierCC resolution

All isolates from badgers were in HierCC cluster HC400_299 (Fig. 3B, dashed box), which also included other isolates from humans and other animals. HC400_299 was investigated by Maximum-Likelihood trees of core, non-repetitive SNPs called against a reference draft genome with the help of the EnteroBase Dendrogram GUI. One tree encompassed 149 isolates from the British Isles which were in EnteroBase prior to establishing the Agama Study group. A second tree (Fig. 4B) contained the final data set of 213 genomes, including isolates from additional badgers and multiple countries. A comparison of the two trees is highly instructive on the effects of sample bias.

**Figure 4.**
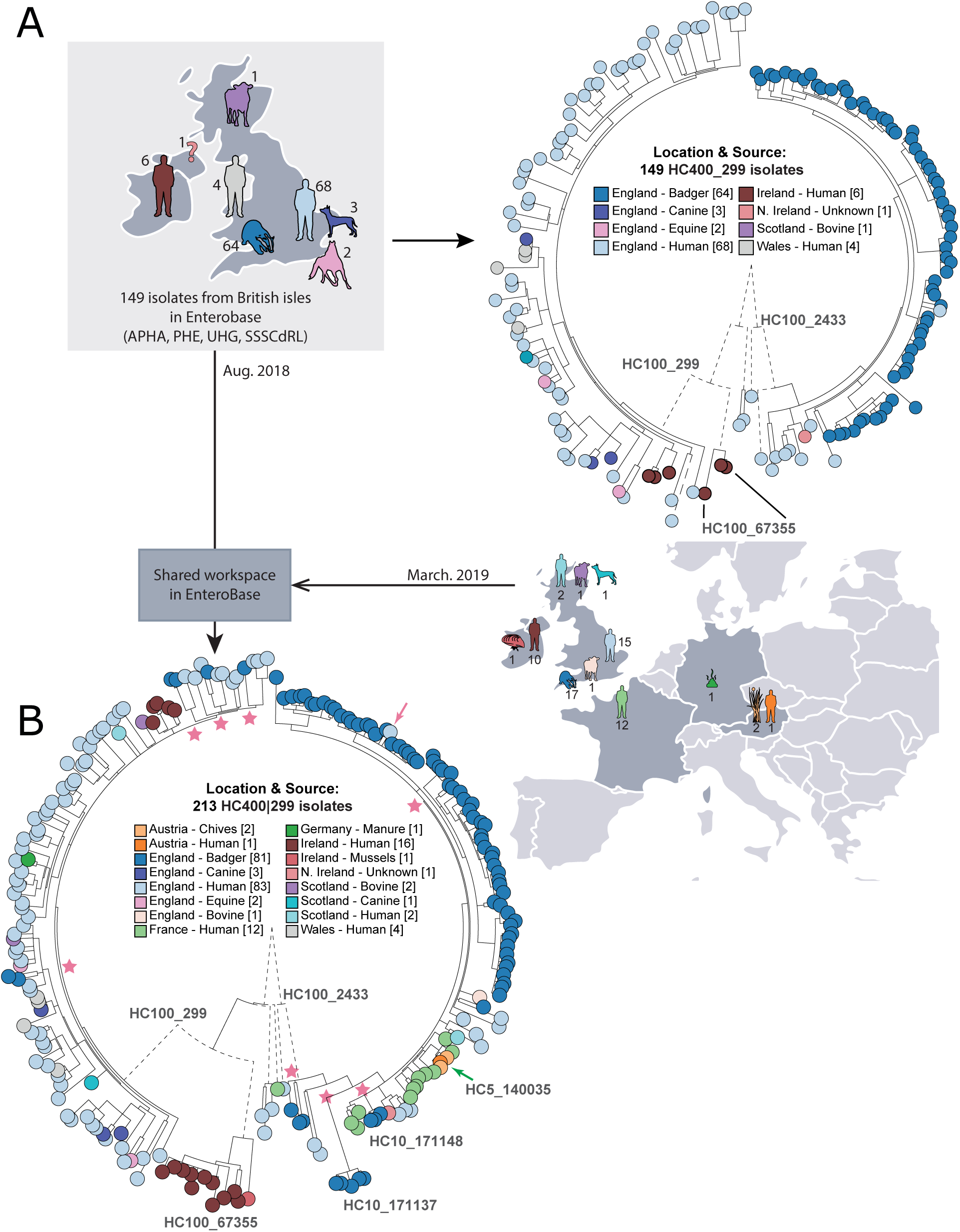
Effects of sample bias on inferred transmission chains within HC400_299 Agama isolates. A) Left: map of hosts in the British Isles of 149 Agama isolates in EnteroBase in August, 2018. Right: Maximum-likelihood radial phylogeny (https://tinyurl.com/AgamaFig4A) based on RAxML (Stamatakis 2014) of 8,791 non-repetitive core SNPs as calculated by EnteroBase Dendrogram against reference genome 283179. Color-coding is according to a User-defined Field (Location & Source). HC100 cluster designations for three micro-clades are indicated. HC100_2433 contained all Agama from badgers. B) Right: summary of hosts and countries from which 64 additional Agama isolates had been sequenced by March 2019. Left: Maximum-likelihood radial dendrogram (https://tinyurl.com/AgamaF4B) based on 9,701 SNPs from 213 isolates. Multiple isolates of Agama in HC100_2433 were now from humans and food in France and Austria. HC100_299 and HC100_67355 now contained multiple isolates from badgers, livestock, companion animals and mussels, demonstrating that the prior strong association of Agama with humans and badgers in part A reflected sample bias. Stars indicate multiple MRCAs of Agama in English badgers while the pink arrow indicates a potential transmission from badgers to a human in Bath/North East Somerset, which is close to Woodchester Park. The green arrow indicates a potential food-borne transmission chain consisting of four closely related Agama isolates in HC5_140035 from Austria (chives x 2; human blood culture x 1) and France (human x 1) that were isolated in 2018. The geographical locations of the badger isolates are shown in Fig. S5.

Almost all of the initial HC400_299 genomes fell into three micro-clades designated HC100_299, HC100_2433 and HC100_67355. All badger isolates were from Woodchester Park (2006-2007) within the context of a long-term live capture-mark-recapture study (McDonald *et al.* 2018). The Agama isolates from those badgers formed a monophyletic clade within HC100_2433, whose basal nodes represented human isolates. This branch topology suggested that a single recent common ancestor of all badger isolates which had been transmitted from humans or their waste products.

The badgers in Woodchester Park occupy adjacent social group territories which each contain several setts (burrows). HC100_2433 contains multiple HC10 clusters of Agama from badgers (Supplemental Fig. S5A). To investigate whether these micro-clusters might mark transmission chains between setts and social groups, a Newick sub-tree of HC100_2433 plus geographical co-ordinates was transmitted from GrapeTree to MicroReact (Argimon *et al.* 2016), an external program which is specialized in depicting geographical associations. Badgers occasionally move between neighboring social groups (Rogers *et al.* 1998). Transmissions associated with such moves are supported by the observation that five distinct HC10 clusters each contained isolates from two social groups in close proximity (Fig. S5B).

#### Long-term dispersals and inter-host transmissions

The 63 additional HC400_299 Agama genomes that were sequenced by the Agama Study Group provided important insights on the dissemination of Agama over a longer time frame, and demonstrated the dramatic effects of sample bias. Seventeen Agama strains had been isolated from English badgers at multiple locations in south-west England between 1998 and 2016 (Fig. S5B), and stored at APHA. Eleven of them were in HC100_2433. However, rather than being interspersed among the initial genomes from badgers, they defined novel micro-clusters, including HC10_171137 and HC10_171148, which were the most basal clades in HC100_2433 (Fig. 4B). The other six badger isolates from additional geographical sources were interspersed among human isolates in HC100_299 (Fig. 4B), which had previously not included any badger isolates (Fig. S5F). These results show that the diversity of Agama from English badgers is comparable to their diversity within English humans, and that it would be difficult to reliably infer the original host of these clades or the directionality of inter-host transmissions. Further observations on micro-epidemiology of Agama transmissions between hosts and countries are presented in Supplemental Material.

### Case Study 2: Combining modern *Y. pestis* genomes with ancient metagenomes

EnteroBase automatically scours sequence read archives for Illumina short reads from cultivated isolates, assembles their genomes and publishes draft assemblies that pass quality control. In October 2019, EnteroBase had assembled >1,300 genomes of *Y. pestis*, including genomes that had already been assigned to population groups (Cui *et al.* 2013), other recently sequenced genomes from central Asia (Eroshenko *et al.* 2017; Kutyrev *et al.* 2018) and numerous unpublished genomes from Madagascar and Brazil. EnteroBase does not upload assembled genomes, for which adequate, automated quality control measures would be difficult to implement. However, EnteroBase administrators can upload such genomes after *ad hoc* assessment of sequence quality, and EnteroBase contains standard complete genomes such as CO92 (Parkhill *et al.* 2001) and other genomes used to derive the *Y. pestis* phylogeny (Morelli *et al.* 2010).

Enterobase also does not automatically assemble genomes from metagenomes containing mixed reads from multiple taxa, but similar to complete genomes, administrators can upload reconstructed ancient genomes derived from SNP calls against a reference genome.

#### Ancient *Y. pestis*

The number of publications describing ancient *Y. pestis* genomes has increased dramatically over the last few years as ancient plague has been progressively deciphered (Bos *et al.* 2011; Wagner *et al.* 2014; Rasmussen *et al.* 2015; Bos *et al.* 2016; Feldman *et al.* 2016; Spyrou *et al.* 2016; Spyrou *et al.* 2018; Margaryan *et al.* 2018; Namouchi *et al.* 2018; Keller *et al.* 2019; Spyrou *et al.* 2019). The metagenomic short reads used to reconstruct these genomes are routinely deposited in the public domain but the reconstructed ancient genomes are not. This practice has made it difficult for non-bioinformaticians to evaluate the relationships between ancient and modern genomes from *Y. pestis*. However, EnteroBase now provides a solution to this problem.

The EnteroBase EToKi calculation package can reconstruct an ancient genome assembly by unmasking individual nucleotides in a fully masked reference genome based on reliable SNP calls from metagenomic data (Supplemental Fig. S6). We ran EToKi on 56 published ancient metagenomes containing *Y. pestis* and the resulting assemblies and metadata were uploaded to EnteroBase. EnteroBase users can now include those ancient genomes together with other reconstructed genomes and modern genomic assemblies in a workspace of their choice, and use the EnteroBase SNP dendrogram module to calculate and visualize a Maximum Likelihood tree (of up to a current maximum of 200 genomes).

Fig. 5 presents a detailed overview of the genomic relationships of all known *Y. pestis* populations from pandemic plague over the last 5,500 years, including 100s of unpublished modern genomes. This tree was manually annotated using a User-defined Field and Custom View with population designations from the literature on modern isolates to include reconstructed ancient genomes. These population designations have now been updated for additional modern genomes from central Asia and elsewhere. An interactive version of this tree and all related metadata in EnteroBase is publicly available (https://tinyurl.com/YpestisSNP), thus enabling its detailed interrogation by a broad audience from multiple disciplines (Green 2018), and providing a common language for scientific discourse.

**Figure 5.**
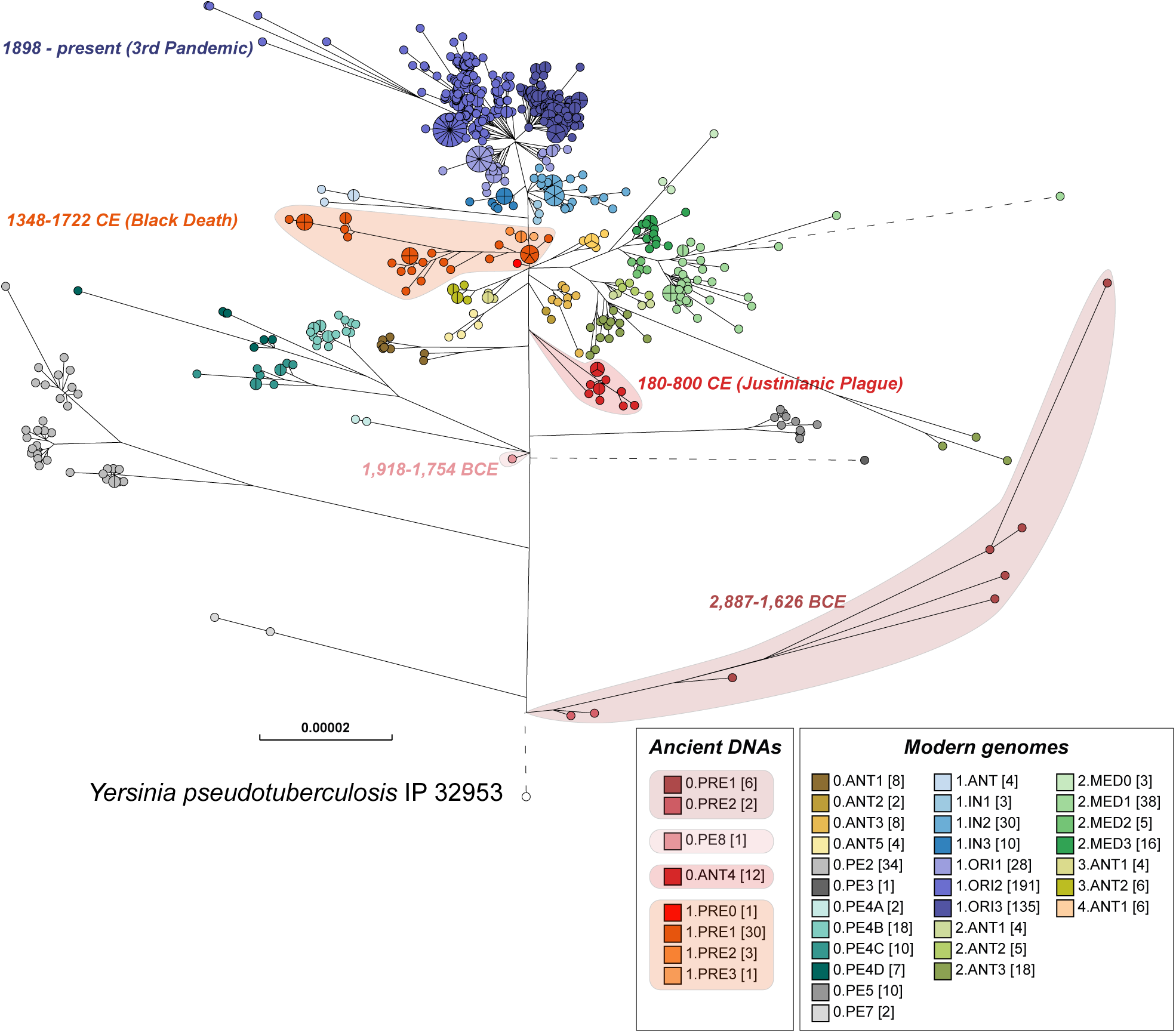
Maximum-Likelihood tree of modern and ancient genomes of *Y. pestis*. EnteroBase contained 1,368 ancient and modern *Yersinia pestis* genomes in October 2019, of which several hundred genomes that had been isolated in Madagascar and Brazil over short time periods demonstrated very low levels of genomic diversity. In order to reduce this sample bias, the dataset used for analysis included only one random representative from each HC0 group from those two countries, leaving a total of 622 modern *Y. pestis* genomes. 56 ancient genomes of *Y. pestis* from existing publications were assembled with EToKi (see Methods), resulting in a total of 678 *Y. pestis* genomes plus *Yersinia pseudotuberculosis* IP32953 as an outgroup (https://tinyurl.com/YpestisWS). The EnteroBase pipelines (Fig. S2D) were used to create a SNP project in which all genomes were aligned against CO92 (2001) using LASTAL. The SNP project identified 23,134 non-repetitive SNPs plus 7,534 short inserts/deletions over 3.8 Mbps of core genomic sites which had been called in ≥95% of the genomes. In this figure, nodes are color-coded by population designations for *Y. pestis* according to published sources (Morelli *et al*. 2010; Cui *et al*. 2013; Achtman 2016), except for 0.PE8 which was assigned to a genome from 1,918-1754 BCE (Spyrou *et al*. 2018). The designation 0.ANT was applied to *Y. pestis* from the Justinianic plague by Wagner et al. 2014, and that designation was also used for a genome associated with the Justinianic plague (DA101) that was later described by Damgaard *et al*., 2018 as 0.PE5.

### Case Study 3: Thinking big – an overview of the core genomic diversity of *Escherichia/Shigella*

*Escherichia coli* has long been one of the primary work-horses of molecular biology. Most studies of *Escherichia* have concentrated on a few well-characterized strains of *E. coli*, but the genus *Escherichia* includes other species: *E. fergusonii, E. albertii, E. marmotae* (Liu *et al.* 2015) and *E. ruysiae* (van der Putten *et al.* 2019). *E. coli* itself includes the genus *Shigella* (Pupo *et al.* 2000), which was assigned a distinctive genus name because it causes dysentery. Initial analyses of the phylogenetic structure of *E. coli* identified multiple deep branches, called haplogroups (Selander *et al.* 1987), and defined the EcoR collection (Ochman and Selander 1984), a classical group of 72 bacterial strains that represented the genetic diversity found with multilocus enzyme electrophoresis. The later isolation of environmental isolates from lakes revealed the existence of “cryptic clades” I-VI which were distinct from the main *E. coli* haplogroups and the other *Escherichia* species (Walk et al. 2009; Luo et al. 2011). Currently, bacterial isolates are routinely assigned to haplogroups or clades by PCR tests for the presence of variably present genes from the accessory genome (Clermont *et al.* 2013) or by programs that identify the presence of those genes in genomic sequences (Beghain *et al.* 2018; Waters *et al.* 2018).

An alternative scheme for subdividing *Escherichia* was introduced in 2006, legacy MLST which includes the assignment of STs to ST Complexes (Wirth *et al.* 2006). Several ST Complexes are common causes of invasive disease in humans and animals, such as ST131 (Stoesser *et al.* 2016; Liu *et al.* 2018), ST95 Complex (Wirth *et al.* 2006; Gordon *et al.* 2017) and ST11 Complex (O157:H7) (Eppinger *et al.* 2011a; Eppinger *et al.* 2011b; Newell and La Ragione 2018). The large number of *Escherichia* genomes in EnteroBase (Table 1) now provides an opportunity to re-investigate the population structure of *Escherichia* on the basis of the greater resolution provided by cgMLST, and within the context of a much larger and more comprehensive sample. in 2018 EnteroBase contained 52,876 genomes. In order to render this sample amenable to calculating an ML tree of core SNPs, we selected a representative sample consisting of one genome from each of the 9,479 *Escherichia* rSTs. In homage to the EcoR collection, we designate this as the EcoRPlus collection.

#### Core genome genetic diversity within *Escherichia*

Homologous recombination is widespread within *E. coli* (Wirth *et al.* 2006). We therefore anticipated that a phylogenetic tree of core genomic differences in EcoRPlus would be ‘fuzzy’, and that ST Complexes and other genetic populations would be only poorly delineated. Instead, considerable core genome population structure is visually apparent in a RapidNJ tree based on pairwise differences at cgMLST alleles between the EcoRPlus genomes (Fig. 6). The most predominant, discrete sets of node clusters were also largely uniform according to cgMLST HC1100 hierarchical clustering. Furthermore, with occasional exceptions, assignments to HC1100 clustering were also largely congruent with ST Complexes based on legacy 7-gene MLST (Supplemental Fig. S7) and with Clermont typing (Supplemental Fig. S8; Supplemental material).

**Figure 6.**
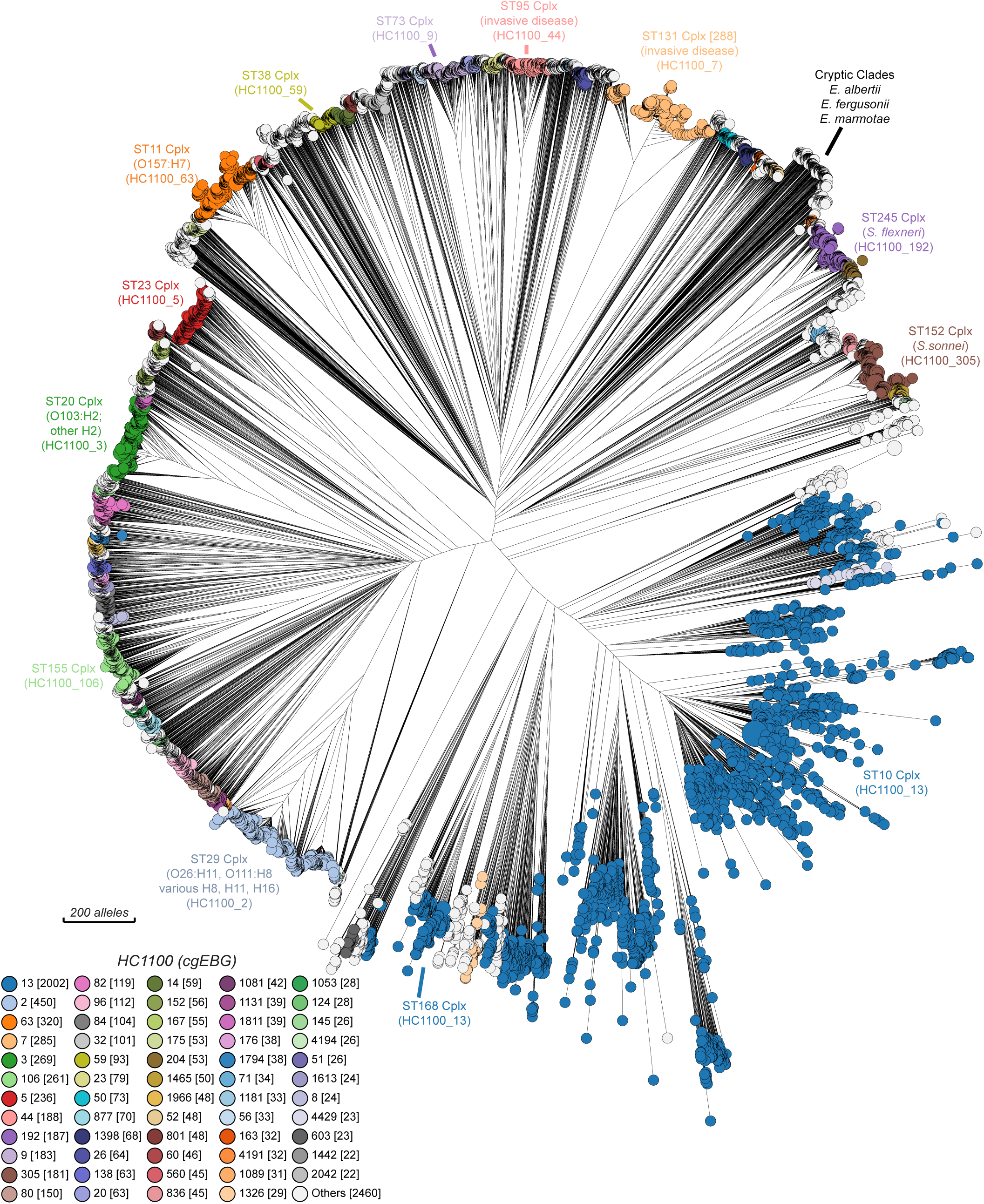
Neighbour-Joining (RapidNJ) tree of core-genome allelic distances in the EcoRPlus Collection of 9,479 genomes. EcoRPlus includes the draft genome with the greatest N50 value from each of the 9,479 rSTs among the 52,876 genomes of *Escherichia* within EnteroBase (August, 2018) (http://tinyurl.com/ECOR-Plus). The nodes in this tree are color-coded by HC1100 clusters, as indicated in the Key at the bottom left. Common HC1100 clusters (plus corresponding ST Complexes) are indicated at the circumference of the tree. These are largely congruent, except that HC1100_13 corresponds to ST10 Complex plus ST168 Complex, and other discrepancies exist among the smaller, unlabeled populations. See Figs S7 and S8, respectively, for color-coding by ST Complex and Clermont typing. An interactive version in which the nodes can be freely color-coded by all available metadata is available at http://tinyurl.com/ECOR-RNJ. A Maximum-Likelihood tree based on SNP differences can be found in Fig. S9.

Fig. 6 may represent the first detailed overview of the entire genetic diversity of the core genome of *Escherichia*. Real time examination of its features (http://tinyurl.com/ECOR-RNJ) is feasible because the GrapeTree algorithm can handle large numbers of cgSTs (Zhou *et al.* 2018a). Nodes can be readily colored by metadata or experimental data (Supplemental Figs. S7-S9), and GrapeTree also readily supports analyses of sub-trees in greater detail. However, although cgMLST allelic distances are reliable indicators of population structures, SNPs are preferable for examining genetic distances. We therefore calculated a Maximum-Likelihood (ML) tree of the 1,230,995 core SNPs within all 9,479 genomes (Supplemental Fig. S9). This tree confirmed the clustering of the members of HC1100 groups within *E. coli*, and also showed that the other *Escherichia* species and cryptic clades II to VIII formed distinct long branches of comparable lengths (Fig. S9 inset).

## Discussion

EnteroBase was originally developed as a genome-based successor to the legacy MLST websites for *Escherichia* (Wirth et al. 2006), *Salmonella* (Achtman et al. 2012), *Yersinia pseudotuberculosis* (Laukkanen-Ninios et al. 2011) and *Moraxella catarrhalis* (Wirth et al. 2007). Its underlying infrastructure is sufficiently generic that EnteroBase was readily extended to *Clostridioides, Helicobacter* and *Vibrio*, and could in principle be extended to other taxa.

EnteroBase was intended to provide a uniform and reliable pipeline that can assemble consistent draft genomes from the numerous short read sequences in public databases (Achtman and Zhou 2014), and to link those assemblies with metadata and genotype predictions. It was designed to provide access to an unprecedentedly large global set of draft genomes to users at both extremes of the spectrum of informatics skills. A further goal was to provide analytical tools, such as GrapeTree (Zhou *et al.* 2018a), that could adequately deal with cgMLST from >100,000 genomes, and Dendrogram, which generates phylograms from non-repetitive core SNPs called against a reference genome. Still another important goal was to support private analyses by groups of colleagues, with the option of subsequently making those analyses publicly available. Case Study 1 illustrates how EnteroBase can be used for all of these tasks and more.

EnteroBase has expanded beyond its original goals, and is morphing in novel directions. It has implemented HierCC for cgMLST, which supports the automated recognition of population structures at multiple levels of resolution (Case Study 1), and may help with the annotation of clusters within phylogenetic trees (Case Study 2; see below). EnteroBase has also been extended to support analyses of metagenomic data from ancient genomes (Zhou *et al.* 2018c) by implementing a subset of the functionality of SPARSE (Zhou *et al.* 2018b) within the stand-alone EToKi package. Case Study 2 illustrates this capability for *Y. pestis.* Additional EnteroBase databases are under development for ancient and modern genomes of *S. enterica* and biofilms within dental calculus. EnteroBase has also demonstrated its capacities for providing overviews of the core genome diversity of entire genera, with currently extant examples consisting of *Salmonella* (Alikhan et al. 2018) and *Escherichia* (Case Study 3).

EnteroBase is already being used by the community to identify genetically related groups of isolates (Johnson *et al.* 2019; Haley *et al.* 2019; Numberger *et al.* 2019; Diemert and Yan 2019), and HierCC has been used to mark international outbreaks of *S. enterica* serovar Poona (Jones *et al.* 2019b) and *E. coli* O26 (Jones *et al.* 2019a). Case Study 1 illustrates how to explore HierCC genomic relationships at multiple levels, ranging from HC2000 (super-lineages) for inter-continental dispersion down to HC5-10 for detecting local transmission chains.

Case Study 1 confirms that although *S. enterica* serovar Agama is rare, it has been isolated from multiple hosts and countries, and is clearly not harmless for humans. The results also document that an enormous sample bias exists in current genomic databases because they largely represent isolates that are relevant to human disease from a limited number of geographic locations.

Case Study 1 may also become a paradigm for identifying long-distance chains of transmission between humans or between humans and their companion or domesticated animals: Four Agama isolates in the HC5_140035 cluster from France (human) and Austria (frozen chives and a human blood culture) differed by no more than 5 of the 3,002 cgMLST loci. These isolates also differed by no more than 5 non-repetitive core SNPs. We anticipate that large numbers of such previously silent transmission chains will be revealed as EnteroBase is used more extensively.

Case study 2 illustrates how EnteroBase can facilitate combining reconstructed genomes from metagenomic sequences with draft genomes from cultured strains. In this case, the metagenomes were from ancient tooth pulp which had been enriched for *Y. pestis*, and the bacterial isolates were modern *Y. pestis* from a variety of global sources since 1898. The resulting phylogenetic tree (Fig. 5) presents a unique overview of the core genomic diversity over 5,000 years of evolution and pandemic spread of plague, which can now be evaluated and used by a broad audience. This tree will be updated at regular intervals as additional genomes or metagenomes become available.

The manual population designations in Fig. 5 are largely reflected by HC10 clusters. However, it is uncertain whether the current HierCC clusters would be stable with time because they were based on only 1,300 *Y. pestis* genomes. EnteroBase will therefore maintain manual annotations in parallel with automated HierCC assignments until a future date when a qualified choice is possible.

Case study 3 defines the EcoRPlus Collection of 9,479 genomes which represents the genetic diversity of 52,876 genomes. It is a worthy successor of EcoR (Ochman and Selander 1984), which contained 72 representatives of 2,600 *E. coli* strains that had been tested by multilocus enzyme electrophoresis in the early 1980s. The genomic assemblies and known metadata of EcoRPlus are publicly available (http://tinyurl.com/ECOR-Plus), and can serve as a reference set of genomes for future analyses with other methods.

Visual examination of an NJ tree of cgMLST allelic diversity color-coded by HierCC HC1100 immediately revealed several discrete *E. coli* populations that have each been the topics of multiple publications (Fig. 6). These included a primary cause of hemolytic uremic syndrome (O157:H7), a common cause of invasive disease in the elderly (the ST131 Complex), as well as multiple distinct clusters of *Shigella* that cause dysentery. However, it also contains multiple other discrete clusters of *E. coli* that are apparently also common causes of global disease in humans and animals but which have not yet received comparable attention. The annotation of this tree would therefore be a laudable task for the entire scientific community interested in *Escherichia.* We also note that HierCC is apparently a one stop, complete replacement for haplogroups, Clermont Typing and ST Complexes, some of whose deficiencies are also illustrated here.

This user’s guide provides an overview of what EnteroBase can do now. With time, we hope to include additional, currently missing features, such as community annotation of the properties of bacterial populations, predicting antimicrobial resistance/sensitivity, and distributing core pipelines to multiple mirror sites. However, EnteroBase is already able to help a broad community of users with a multitude of tasks for the selected genera it supports. More detailed instructions are available in the online documentation (https://enterobase.readthedocs.io/en/latest/) and questions can be addressed to the support team.

## Methods

### Isolation of serovar Agama from badgers

Supplemental Fig. S5B provides a geographical overview of the area in Woodchester, Gloucestershire in which badger setts and social groups were investigated in 2006-2007. This area has been subject to a multi-decade investigation of badger mobility and patterns of infection with *Mycobacterium bovis* (McDonald *et al.* 2018). According to the standard protocol for that study, badgers were subjected to routine capture using steel mesh box traps baited with peanuts, examination under anesthesia and subsequent release. Fecal samples were cultivated at University of Liverpool after selective enrichment (Rappaport-Vassiliadis broth and semi-solid agar), followed by cultivation on MacConkey agar. Lactose-negative colonies that swarmed on Rappaport-Vassiliadis agar but not on nutrient agar, and were catalase-positive and oxidase-negative, were serotyped by slide agglutination tests according to the Kaufmann and White scheme. Additional isolates from badgers from the geographical areas in England that are indicated in Fig. S5D and S5F were collected during routine investigations of animal disease at the APHA.

### Laboratory manipulations and genomic sequencing

At University of Warwick, *Salmonella* were cultivated and DNA was purified by automated procedures as described (O’Farrell et al. 2012). Paired-end 150 bp genomic sequencing was performed in multiplexes of 96-192 samples on an Illumina NextSeq 500 using the High Output Kit v2.5 (FC-404-2002) according to the manufacturer’s instructions. Other institutions used their own standard procedures. Metadata and features of all 344 genomes in Fig. 4 are publicly available in EnteroBase in the workspace ‘Zhou et al. All Agama strains’ (https://tinyurl.com/AgamaWS).

### Integration of ancient *Yersinia pestis* genomes in EnteroBase

Metagenomic reads from ancient samples may contain a mixture of sequence reads from the species of interest as well as from genetically similar taxa that represent environmental contamination. In order to deal with this issue and remove such non-specific reads after extraction with the EToKi prepare module, the EToKi assemble module can be used to align the extracted reads after comparisons with an ingroup of genomes related to the species of interest and with an outgroup of genomes from other species. In the case of Fig. 5, the ingroup consisted of *Y. pestis* genomes CO92 (2001), Pestoides F, KIM10+ and 91001 and the outgroup consisted of genomes *Y. pseudotuberculosis* IP32953 and IP31758, *Y. similis* 228 and *Y. enterocolitica* 8081. Reads were excluded which had higher alignment scores to the outgroup genomes than to the ingroup genomes. Prior to mapping reads to the *Y. pestis* reference genome (CO92 (2001)), a pseudo-genome was created in which all nucleotides were masked in order to ensure that only nucleotides supported by metagenomic reads would be used for phylogenetic analysis. For the 13 ancient genomes whose publications included complete SNP lists, we unmasked the sites in the pseudo-genomes which were included in the published SNP lists. For the other 43 genomes, the filtered metagenomic reads were mapped onto the pseudo-genome with minimap2 (Li 2018), and evaluated with Pilon (Walker *et al.* 2014), and sites in the pseudo-genome were unmasked which were covered by ≥3 reads and had a consensus base that was supported by ≥80% of the mapped reads. All 56 pseudo-genomes were stored in EnteroBase together with their associated metadata.

## Data Access

The Illumina sequence reads for 161 new genomes of *S. enterica* serovar Agama generated in this study have been submitted to the European Nucleotide Archive database (ENA; https://www.ebi.ac.uk/ena) under study accession numbers ERP114376, ERP114456, ERP114871 and ERP115055. The genomic properties, metadata and accession codes for the 329 genomic assemblies in HC2000_299 are summarized in Supplemental Table S3 and in Online Table 1 (https://wrap.warwick.ac.uk/128112). The metadata, genomic assemblies and annotations are also available from the publicly available workspace “Zhou et al. All Agama Strains” (https://tinyurl.com/AgamaWS). The EToKi package and its documentation are accessible at https://github.com/zheminzhou/EToKi. EnteroBase documentation is accessible at https://enterobase.readthedocs.io/en/latest/. An interactive version of Figure 3 is available at https://tinyurl.com/AgamaFig3. Trees presented in Fig. 4 are available separately at (A) https://tinyurl.com/AgamaFig4A and (B) https://tinyurl.com/AgamaF4B. An interactive version of Fig. 5 is available at https://tinyurl.com/YpestisSNP. The MicroReact projects of Figure S5 are available at (A,B) https://microreact.org/project/t7qlSSslh/3e634888; (C,D) https://microreact.org/project/9XUC7i-Fm/fed65ff5 and (E,F) https://microreact.org/project/XaJm1cNjY/69748fe3. The tree shown in Figure 6 as well as Supplemental Figs. S7-S8 are available at http://tinyurl.com/ECOR-RNJ;

## Acknowledgements

EnteroBase development was funded by the BBSRC (BB/L020319/1) and the Wellcome Trust (202792/Z/16/Z). We gratefully acknowledge sharing of strains and data by Niall Delappe and Martin Cormican, Salmonella Reference Laboratory, Galway, Ireland, and critical comments on the text by Nina Luhmann and Jane Charlesworth.

## Author Contributions

MA wrote the manuscript with the help of all other authors. ZZ, N-FA, KM and YF were responsible for the development of EnteroBase under the guidance of MA. N-FA and KM were responsible for the online manual. Analyses were performed and figures were drawn by ZZ and N-FA under the guidance of MA. The Agama Study Group provided information, bacterial strains, DNAs and genomic sequences from Agama isolates from all over Europe, and was involved in writing the manuscript and evaluating the conclusions.

## Abbreviations

wgMLST: whole genome MultiLocus Sequence Typing (Maiden *et al.* 2013);
cgMLST: core genome MultiLocus Sequence Typing (Mellmann *et al.* 2011);
rMLST: ribosomal MultiLocus Sequence Typing (Jolley *et al.* 2012).

