## Supplemental Material for "The user’s guide to comparative genomics with EnteroBase, including case studies on transmissions of micro-clades of *Salmonella*, the phylogeny of ancient and modern *Yersinia pestis* genomes, and the core genomic diversity of all *Escherichia*"

### EnteroBase Metaparser conversions of metadata for host or environment.

Metadata within short read archives are associated with a wide range of designations for the hosts or environments from which the bacteria were isolated. This raw data was stored in EnteroBase as Source->Details. In 2015, we manually assigned 3,546 distinct Source->Details entries to pre-defined categories of Source->Niche and Source->Type (Supplemental Fig. S3; Supplemental Table S4). 2,000 of these entries provided an initial training set and the remaining 1,546 entries served as a test set for a Native Bayesian source classifier that is implemented in the NLTK Natural Language Toolkit for Python (Bird *et al.* 2009). The NLTK classifier achieved an accuracy of ~80% on the test set after initial training. We then re-trained the source classifier using all 3,546 manually curated entries, and have since used it continuously to assign GenBank metadata into the nested Source metadata fields within EnteroBase.

During curation over the following two years, we found a number of obvious misclassifications. We therefore retested the source classifier in 2018 with an independent set of 3,000 manually curated recent entries. Those tests yielded an accuracy of only 60%, which reflects the large number of novel designations which are currently being used, many of which were not included in the initial training set and are not recognized by the source classifier. When such mistakes are observed during curation, they are corrected. However the number of existing EnteroBase curators is insufficient to support the manual curation of the 100,000s of entries in EnteroBase, and EnteroBase will continue to possess multiple false assignments to Source categories until the source classifier can be updated.

### An example of using Uberstrains and sub-strains.

The EnteroBase *Yersinia* database includes two distinct genomes for *Y. pestis* CO92, the genome originally sequenced in 2001 (Parkhill et al. 2001) and a subsequent, corrected genome from 2015 (Johnson et al. 2015). The more recent genome sequence is the Uberstrain for genomes from multiple bacterial colonies that were sequenced after infecting prairie dogs in the laboratory with CO92 (project PRJNA340278) (Supplemental Fig. S4). The older genome is maintained in

EnteroBase as a separate Uberstrain because it was previously used as a reference genome for SNP calling (Morelli et al. 2010; Bos et al. 2011), and continues to be used as a reference for *Y. pestis* ancient DNA (aDNA) projects based on archaeological samples (Wagner et al. 2014; Rasmussen et al. 2015; Bos et al. 2016; Feldman et al. 2016; Spyrou et al. 2016; Spyrou et al. 2018; Margaryan et al. 2018). Fig. S4 illustrates these points, as well as showing how to load and display sub-strains as well as Uberstrains,

**Micro-epidemiology of Agama transmission between humans and countries.**

The additional Agama genomes provided by the Agama Study Group included isolates from humans in Ireland, France or Austria, as well as multiple isolates from animals and food. The additional genomes showed that HC100_299 was also isolated from humans in Ireland as well as from dogs, cows and horses. (Fig. 4B). All isolates in HC100_67355 were from Ireland, consisting of one clade of 12 isolates from humans and a second clade containing one isolate from mussels, HC100_2433 now contained not only isolates from the British Isles, but also isolates from France and Austria. We were particularly struck by four genomes in HC5_140035 (Fig. 4B, green arrow at 04:00), which had all been isolated in 2018. Three of these isolates were from Austria, two from frozen chives and one from a blood culture from a case of human septicemia. The Austrian isolates differ from each other in pair-wise comparisons by 2-5 non-repetitive core SNPs and 2-4 cgMLST loci. The fourth isolate in HC5_140035 was from a human in France. It differed by 5 SNPs and 5 cgMLST loci from the three Austrian isolates and by 8-35 SNPs and 6-23 cgMLST loci from other Agama in France. We do not know of any epidemiological data that support food-borne transmission of these organisms between France and Austria, or that indicate that frozen chives are a vehicle for food-borne invasive Agama disease. However, this observation is a strong signal for recent transmission chains of Agama between France and Austria with the possible involvement of frozen food products.

**Comparison of Clermont typing with HC1100 clustering**

EnteroBase provides phylotype predictions for *Escherichia* based on Clermont typing according to two distinct algorithms (Beghain *et al.* 2018; Waters *et al.* 2018). The assignments by both Clermont typing algorithms are largely congruent. And those assignments are also largely congruent with the general structure of the ML tree (Fig. S9 inset). However, Clermont typing is based on the presence/absence of accessory genes, which are more variable than are sequences of the core genome. As a result, close examination of the ML tree shows multiple exceptions, as indicated by flashes of discrepant colors in Fig. S8. Clermont typing never included *S. flexneri* (HC1100_192) and scores it incorrectly as haplogroup A, or *S. sonnei,* which it scores as Unknown (Fig. S8).
