## Supplemental Table S1 for "The user’s guide to comparative genomics with EnteroBase, including case studies on transmissions of micro-clades of *Salmonella*, the phylogeny of ancient and modern *Yersinia pestis* genomes, and the core genomic diversity of all *Escherichia*"

**Supplemental Table S1.** Details of quality controls settings that are used by Enterobase for draft assemblies.

| Genus | min size (KBps) | max size (KBps) | min N50 (KBps) | max number of contigs | max low quality sites | min taxonomic purity |
|---|---|---|---|---|---|---|
| ***Salmonella*** | 4,000 | 5,800 | 20 | 600 | 5% | 70% |
| ***Escherichia/ Shigella*** | 3,700 | 6,400 | 15 | 800 | 5% | 70% |
| ***Yersinia*** | 3,700 | 5,500 | 15 | 600 | 5% | 65% |
| ***Clostridioides*** | 3,600 | 4,800 | 20 | 600 | 5% | 65% |
| ***Moraxella*** | 1,800 | 2,600 | 20 | 600 | 5% | 65% |
| ***Helicobacter*** | 1,300 | 3,000 | 10 | 600 | 5% | 65% |
| ***Vibrio*** | 3,000 | 5,900 | 20 | 700 | 5% | 65% |
