## Supplemental Table S2 for "The user’s guide to comparative genomics with EnteroBase, including case studies on transmissions of micro-clades of *Salmonella*, the phylogeny of ancient and modern *Yersinia pestis* genomes, and the core genomic diversity of all *Escherichia*"

**Supplemental Table S2A.** Metadata fields used in all EnteroBase databases.

| Field | Description | Special features and editing format |
|---|---|---|
| Uberstrain | Primary entry | Place holder for multiple almost identical entries |
| Name | Strain designation | Not necessarily unique |
| Comment | Free text | Unrestricted text |
| Data Source[1] | Properties of short reads | Includes Accession number. Special editing dialog box |
| Source-> Source Niche[2] | Ecological niche | Dropdown list (Human, Aquatic, Food, etc.) |
| Source-> Source Type[2] | Taxonomy of source | Dropdown list (Human, Avian, Camelid, etc.) |
| Source-> Source Details[2] | Free Text | Additional details |
| Collection Date-> Year[3] | Year of sample | Text with calendar date sub-dialog |
| Collection Date-> Month[3] | Month of sample | Text with calendar date sub-dialog |
| Collection Date-> Day[3] | Day of sample | Text with calendar date sub-dialog |
| Collection Date-> Time[3] | Time of sample | Text |
| Location-> Continent[4] | Geographical data | Text with browser suggestions. Google map. |
| Location-> Country[4] | Geographical data | Text with browser suggestions. Google map. |
| Location-> Region[4] | Geographical data | Text with browser suggestions. Google map. |
| Location-> District[4] | Geographical data | Text with browser suggestions. Google map. |
| Location-> City[4] | Geographical data | Text with browser suggestions. Google map. |
| Location-> Post Code[4] | Geographical data | Text. Google map |
| Location-> Latitude[4] | Geographical data | Text. Google map. |
| Location-> Longitude[4] | Geographical data | Text. Google map. |
| Lab Contact | Free Text | Institution for contact |
| Species | Species or sub-species | Dropdown list |
| Project-> Bio Project ID[5] | NCBI BioProject Accession | Non-editable URL |
| Project-> Project ID[5] | NCBI SRA Study | Non-editable URL |
| Sample-> Sample ID[6] | NCBI BioSample Accession | Non-editable URL |
| Sample-> Secondary Sample ID[6] | NCBI SRA Secondary Sample | Non-editable URL |
| Date Entered | Initial publication date | Non-editable information |
| Release Date | Future release date (<12 m) | User-specified delay for public downloads |
| Barcode | Internal EnteroBase ID | Unique identifier for each database entry |

NOTES:

[1]Data source has the format (Accession No.;Sequencing Platform;Sequencing Library;Insert Size;Experiment;Status) when downloaded

[2]Composite of multiple sub-fields. To select default field right click on Header “Source”.

[3] Composite of multiple sub-fields. Default field selectable through right click on Header “Collection Date. To edit year, month and day, click on Calendar icon.

[4]Composite of geographical data supported by a Google Maps depiction. Same format as MicroReact. To select default field right click on Header “Location”.

[5]Composite of Project information as stored at GenBank/NCBI. To select default field right click on Header “Project”.

[6]Composite of Sample information as stored at GenBank/NCBI. To select default field right click on Header “Sample”.

**Supplemental Table S2B.** Database-specific fields.

| Field | Description | Special features and editing format |
|---|---|---|
| *Escherichia/Shigella* | Legacy fields from legacy MLST | |
| Serological Group | Common descriptor e.g. K1 | Dropdown list of common names |
| Serotype | Antigenic formula | Text box |
| EcoR Cluster | Haplogroup designation | Dropdown list |
| Path/Nonpath | Pathogen or Non pathogen | Dropdown list |
| Simple Patho | Acronyms for pathovars | Dropdown list |
| Simple Disease | Clinical disease category | Text box |
| Disease | Clinical disease category | Dropdown list |
| *Salmonella* | | |
| Disease | Clinical disease category | Text box |
| Antigenic Formulas | O:H1:H2 formula | Text box |
| Phage Type | Phage typing designation | Text box |
| Serovar | Kaufmann-White Serovar name | Text box with browser suggestions |
| Subspecies | I, II, …VII, *S. bongori*, novel subsp. A-C | Dropdown list |
| *Clostridioides* | | |
| PCR Ribotype | Classical ribotype designation (3-digits) | Text box |
| *Helicobacter* | | |
| Alias | Alternative strain name | Text box |
| Antimicrobial Resistance | List of resistances | Text box |
| Citations | PubMed IDs | Text box |
| Contact | Name and/or Email address of contact person | Text box |
| Source-> Host Age | Age of carrier | Multiline dialog. Text box. |
| Source-> Host Ethnicity | Ethnicity of carrier | Multiline dialog. Text box. |
| Source-> Host Sex | Sex of carrier | Dropdown list |
| Uploader | Lab source of data | Text box |
| *Yersinia* | Legacy fields from legacy MLST | |
| Disease | Clinical disease category | Text box |
| Alternative Name | Alternative strain name | Text box |
| Biogroup | Sub-species based on phenotype, e.g. Orientalis | Text box |
| Contact Name | Laboratory who uploaded reads | Text box |
| O Serotype | Serological formula | Text box |
| *Moraxella* | Legacy fields from legacy MLST | |
| Path/Nonpath | Pathogen or Non pathogen | Dropdown list |
