## Supplemental Table S4 for "The user’s guide to comparative genomics with EnteroBase, including case studies on transmissions of micro-clades of *Salmonella*, the phylogeny of ancient and modern *Yersinia pestis* genomes, and the core genomic diversity of all *Escherichia*"

**Supplemental Table S4.** EnteroBase metaparser classification scheme for Source Niche and Source Type based on Source Details

| Source Niche | Source Type | Examples of Source Details |
|---|---|---|
| Aquatic | Fish; Marine Mammal; Shellfish | Tuna, lobster |
| Companion Animal | Canine; Feline | Cat, dog |
| Environment | Air; Plant; Soil/Dust; Water | River, tree, soil |
| Feed | Animal Feed; Meat | Dog treat, fishmeal |
| Food | Composite Food; Dairy; Fish; Meat; Shellfish | Milk, salami, ready-to-eat food |
| Human | Human | Patient, biopsy |
| Laboratory | Laboratory | Reference strain, serial passage |
| Livestock | Bovine; Camelid; Equine; Ovine; Swine | Horse, calf |
| Poultry | Avian | Turkey, chicken |
| Wild Animal | Amphibian; Avian; Bat; Bovine; Camelid; Canine; Deer; Equine; Feline; Invertebrates; Marsupial; Other Mammal; Ovine; Primate; Reptile; Rodent; Swine | Flamingo, frog, python, Spider |
| ND | ND | |
