## Supplemental Figures S1-S9 for "The user’s guide to comparative genomics with EnteroBase, including case studies on transmissions of micro-clades of *Salmonella*, the phylogeny of ancient and modern *Yersinia pestis* genomes, and the core genomic diversity of all *Escherichia*"

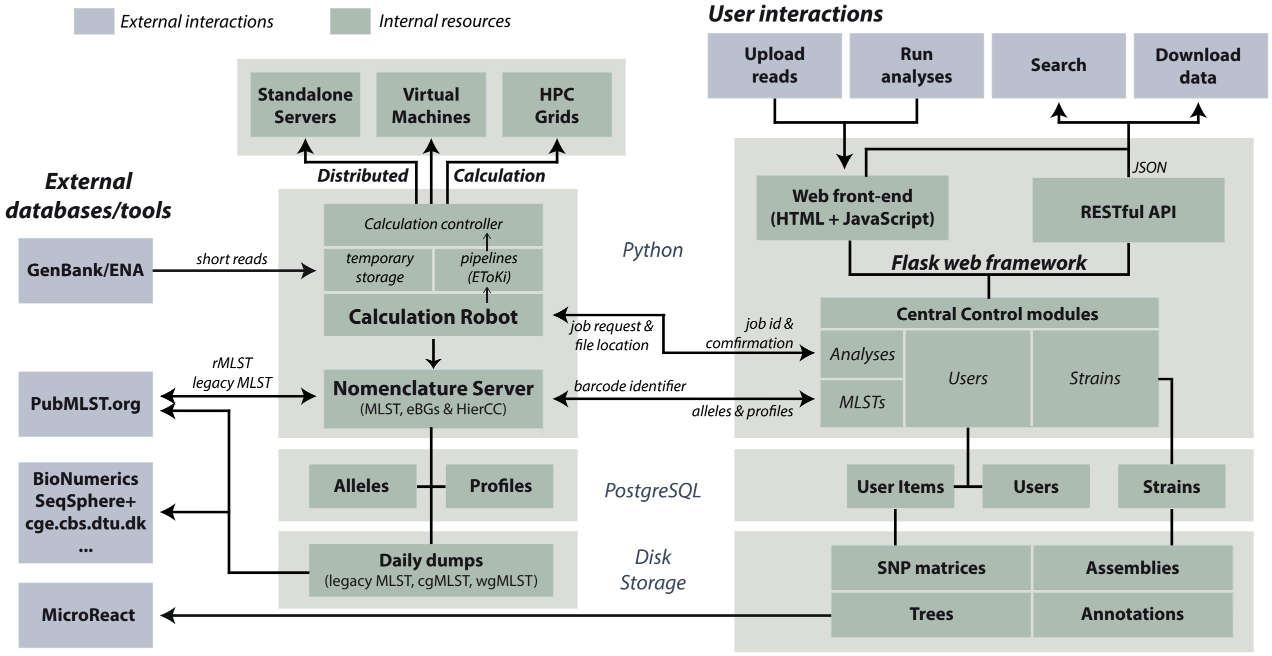


**Supplemental Figure S1. Overview of workflow within EnteroBase.** User interactions (upper right): EnteroBase supports user interactions via a web browser front end or a RESTful API (Application Programming Interface; details at http://tinyurl.com/EnteroBaseAPIDocs), both of which connect to a Python Flask web framework environment that handles all further interactions with structured data (PostgreSQL databases) and stored files in disk storage via Central Control modules. These Central Control modules are also responsible for interacting via JSON strings with the Calculation Robot and Nomenclature Server. The Calculation Robot is responsible for automatically uploading novel short read sequences from GenBank, and uses EToKi (Fig. S2) to assemble and annotate draft genomes, and perform other calculations. The Nomenclature Server (NServ) is responsible for defining and calling MLST alleles and Sequence Types (STs), as well as maintaining population structure designations and assignments. NServ also communicates with external databases; it exchanges data with PubMLST and provides daily dumps of legacy MLST, cgMLST and wgMLST data for external databases and commercial providers.

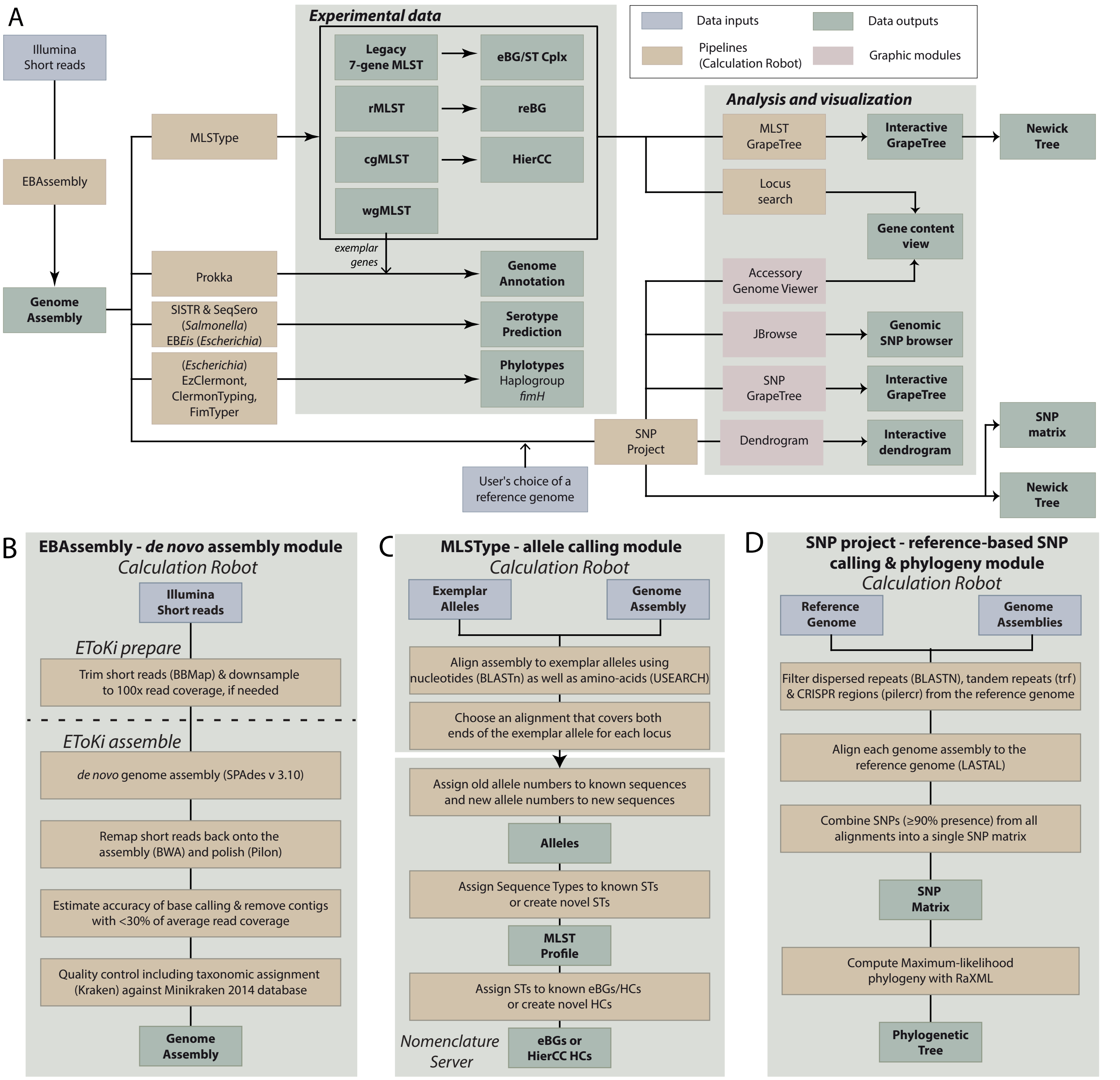


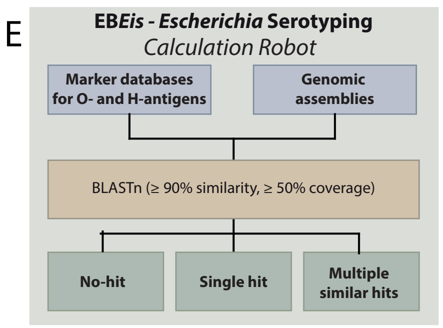


**Supplemental Figure S2. EnteroBase algorithms and the EToKi toolkit.** The backend pipelines within EnteroBase use multiple calculation and graphic modules, to produce a large variety of data and graphical outputs (https://enterobase.readthedocs.io/en/latest//about.html), including a variety of stand-alone programs. A) Algorithm flow. Left: Illumina short reads are assembled in genomes, which are then used for MLST typing at multiple levels of resolution, annotation, and the prediction of serotypes and phylotypes. Right: Tools for the analysis and visualization of genetic distances (MLST alleles or SNPs) and metadata for selected database entries. B-E) Four modules within EToKi (EnteroBase ToolKit), a publicly available standalone package for external command line usage of EnteroBase pipelines that are potentially of general interest. B) EBAssembly performs assemblies, including trimming, automatic down-sampling from enormous datasets, remapping and polishing. EBAssembly estimates the accuracy of base calls and the most probable taxonomic sources of the sequences (Kraken against MiniKraken database (Wood and Salzberg 2014)). These summary statistics are presented in the Experimental Data field "Assembly stats". EBAssembly also provides access to the sub-modules 'prepare' and 'assemble', which can be used for extracting genomic assemblies from metagenomic data (see Supplemental Fig. S6). Standalone programs: BBMap in BBTools (Bushnell 2016), SPADES (Bankevich et al. 2012), BWA (Li and Durbin 2010), Pilon (Walker et al. 2014). C) MLSType calls MLST alleles from genomic assemblies. EnteroBase maintains "Exemplar Alleles", consisting of one allele sequence for each gene in the wgMLST scheme and in legacy MLST. (rMLST and cgMLST are subsets of the wgMLST scheme.) The Exemplar Alleles are aligned to each genome assembly with BLASTn (Altschul et al. 1990) or the USEARCH module UBlastP (Edgar 2010) in order to map the ends of their corresponding loci and extract their sequences. These are assigned to existing allele numbers and STs for each of the MLST schemes unless they are novel, in which case new numbers are assigned. STs are assigned to known eBGs or HC clusters or used to define new clusters. In EnteroBase itself, these tasks are separately performed by the Calculation Robot and the Nomenclature Server. D) An EnteroBase SNP project first masks repetitive dispersed repeats (BLASTN) or tandem repeats (trf (Benson 1999)) or CRISPR regions (pilercr (Edgar 2007)) in the reference genome, and then aligns each genome assembly to that partially masked reference genome (LASTAL in Last (Kielbasa et al. 2011)). The resulting SNP matrix can be downloaded by the user and/or used to calculate an ML phylogeny (RAxML V8 (Stamatakis 2014)). Within EnteroBase, ML phylogenies can be calculated for up to 200 genomes and then displayed by GrapeTree or Dendrogram. SNP Project in EToKi is suitable for larger projects because the number of genomes is not limited except by computer hardware constraints. However, EToKi does not include graphical visualisation tools. E) EBEis predicts Escherichia O serotypes based on the marker genes *wzx, wzy, wzt, and wzm* (Joensen et al. 2015; DebRoy et al. 2016) and H serotypes based on the flagellar *fliC* antigen (Joensen et al. 2015). The standalone version of EToKi including the standalone programs cited above is available at GitHub (https://github.com/zheminzhou/EToKi).

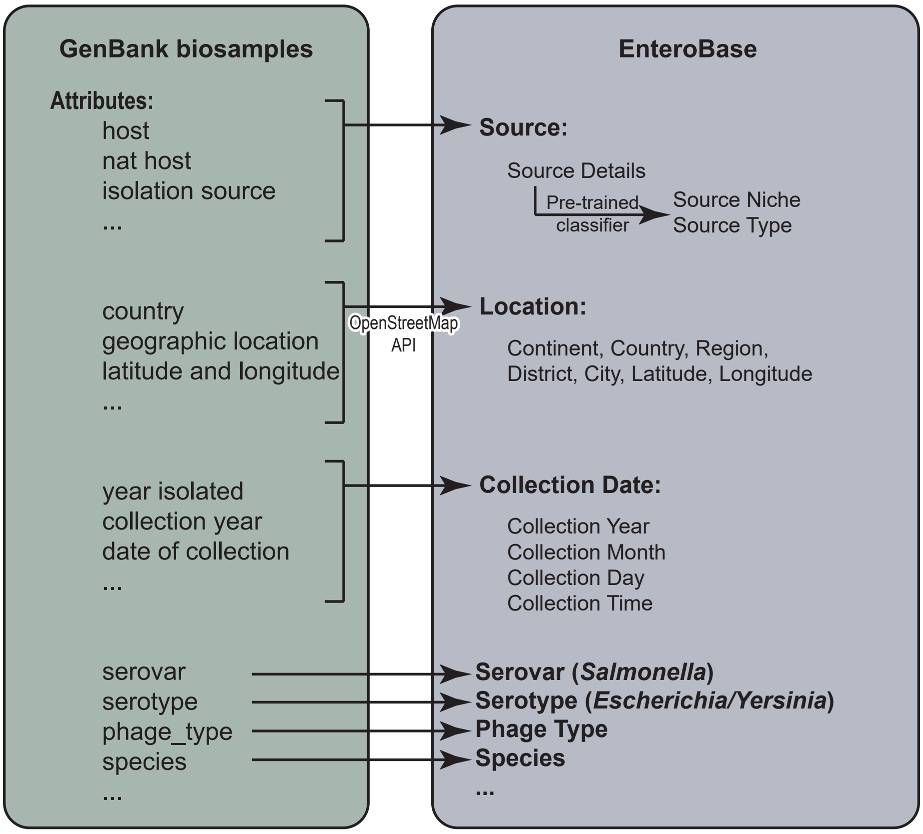


**Supplemental Figure S3. Correspondences between metadata fields in GenBank and in EnteroBase.** EnteroBase metadata fields (Supplemental Table S2-S3) are somewhat different from metadata fields in GenBank. GenBank metadata in the categories Source, Location, Date and Others are therefore automatically converted by an automated metaparser into EnteroBase metadata categories as part of the uploading process. EnteroBase stores the original text from GenBank Source in Source Details and classifies those contents into Source Niche and Source Type (Table S3), all of which are combined within the Source composite field (Table S2A).The other categories of GenBank metadata correspond closely to EnteroBase metadata categories except that some of them are presented as composite fields within EnteroBase (Table S2A).

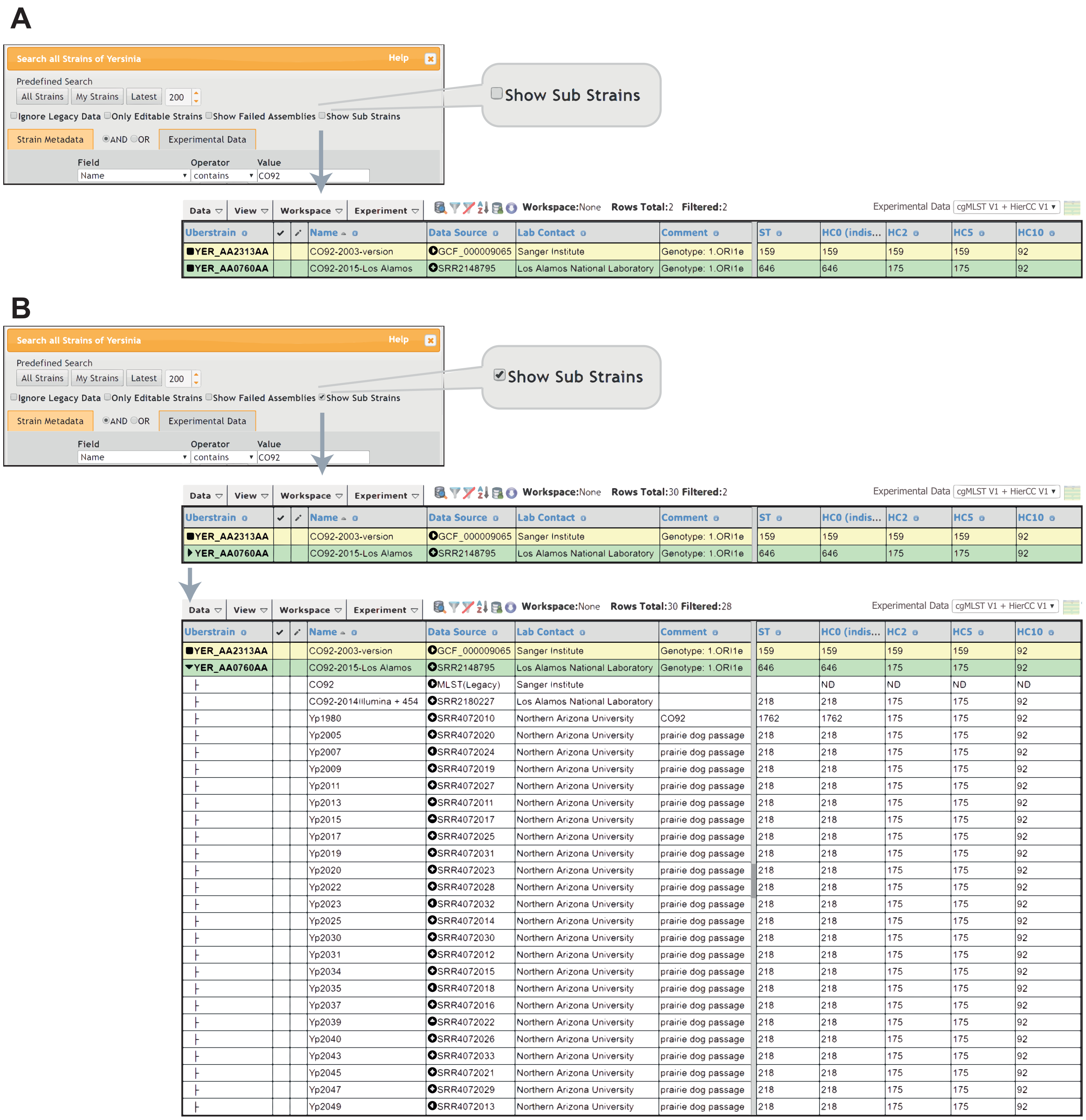


**Supplemental Figure S4. Uberstrains and sub-strains.** A) In its default mode, Show Sub Strains is unticked within the Search Dialog (top), and only Uberstrains are retrieved by the workspace (bottom), as indicated by a black square to the left of the Uberstrain barcode designation at the left. The example shows two distinct Uberstrains of *Y. pestis* CO92, one which was sequenced in 2001 and a second sequenced in 2015, in which 13 erroneous SNP calls have been corrected. B) When Show Sub Strains is checked in the search dialog, the browser shows a triangle at the far left of Uberstrains that contain one or more Sub-Strains. Clicking on that triangle opens a previously hidden tree-like hierarchy containing all its sub-strains. To open these hierarchies for all Uberstrains in the browser window, choose View\Show All Sub-strains in the top browser Menu. View\Close all Sub-strains reverts to showing only Uberstrains.

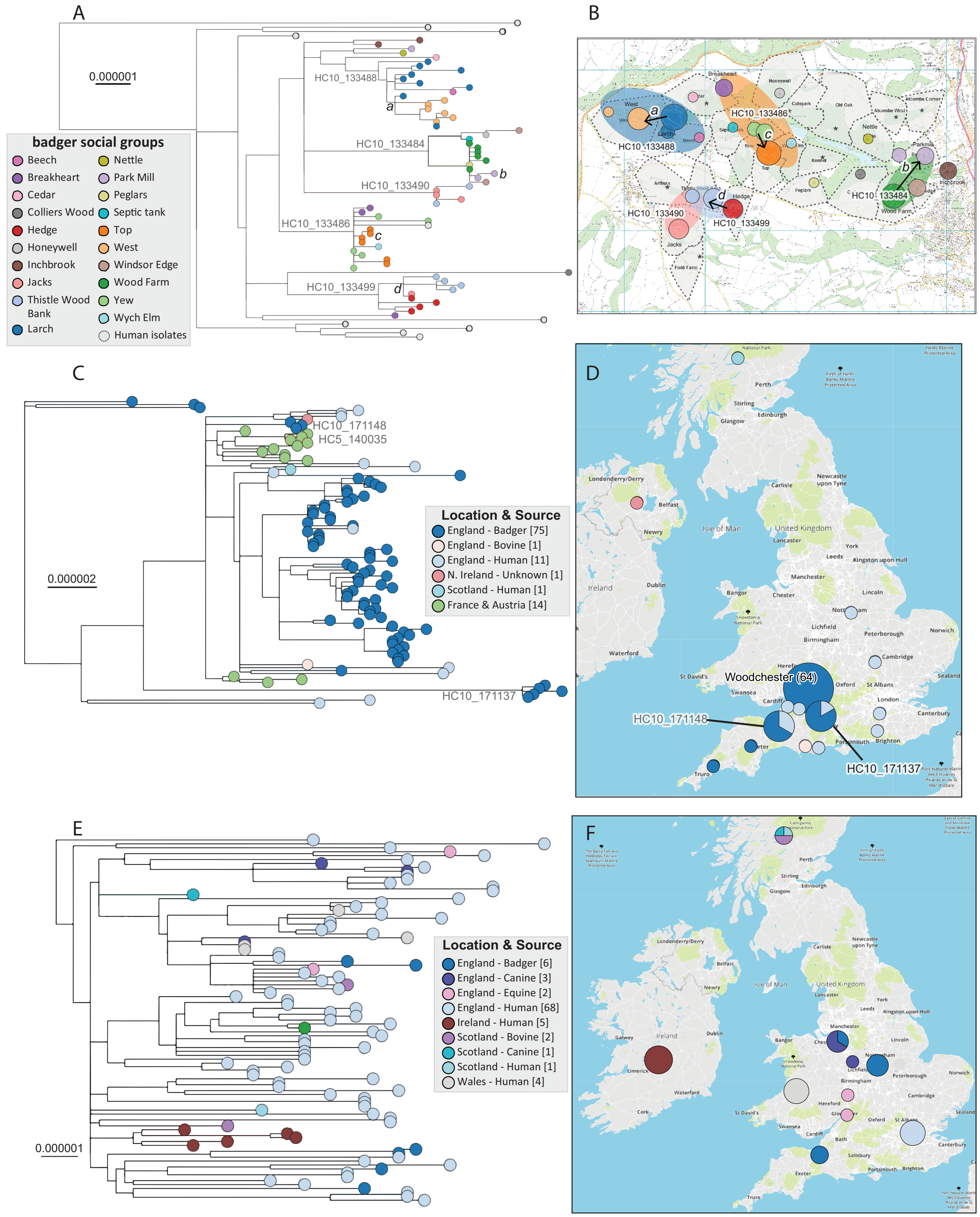


**Supplemental Figure S5. Phylodynamics of isolates from badgers.** GrapeTree was used to transfer individual subtrees plus their GPS coordinates and metadata to MicroReact (Argimon et al. 2016) (http://tinyurl.com/GrapeTreeRefManual). (A, C, E) Phylogenetic trees drawn by MicroReact. (B, D, F) Maps of geographic locations within MicroReact, except that in part B, where the MicroReact tree was overlaid by the idealized spatial distributions of badger social groups and setts as elucidated by (McDonald et al. 2018). (A, B) Sixty four Agama isolates from badgers in Woodchester Park that were collected in 2006-2007 plus 10 related isolates from humans. Five HC10 clusters of genetically related genomes were isolated from neighbouring badger social groups (colored ovals in part B), of which four are inferred to have moved by local transmission chains *a, b, c, d* as indicated in part B (https://microreact.org/project/t7qlSSslh/3e634888). (C, D) 103 Agama isolates in HC100_2433, including 75 from badgers in Woodchester Park and elsewhere in England that were collected between 1998 and 2010 (https://microreact.org/project/9XUC7i-Fm/fed-65ff5). (E, F) 92 Agama isolates in HC100_299 from the British Isles, including 6 from badgers that were collected between 2009 and 2016 (https://microreact.org/project/XaJm1cNjY/69748fe3).

A

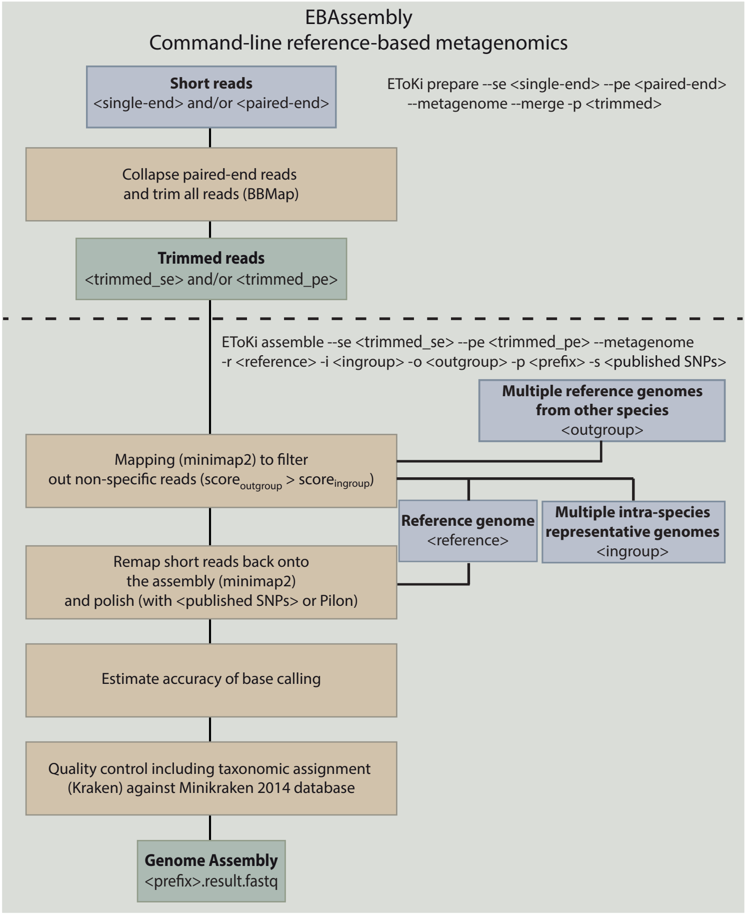


**Supplemental Figure S6. Extracting aDNA assemblies from metagenomic sequences with the EBAssembly module of EToKi.** EBAssembly includes functions for extracting genome-specific reads from metagenomic sequences which are only accessible in the stand-alone, command-line version of EToKi. The EToKi prepare module can collapse paired-end reads and trim both paired-end and single-end reads without down-sampling. As described in the documentation (https://github.com/zheminzhou/EToKi), the EToKi assemble module incorporates elements from SPARSE (Zhou et al. 2018) to identify genome-specific short reads within metagenomic sequences after specifying a reference genome sequence, an in-group of related genomes and a related but distinct out-group of genomes . The module replaces nucleotides in the reference genome by their calculated SNVs after checking that they are supported by at least 3 metagenomic reads, and the supporting read frequencies occur with at least one-third of the average read depth. It also allows constraining SNP calls to (published) SNPS within a text file, and save the modified sequence of the reference genome in a form which can be uploaded to EnteroBase by admins and curators.

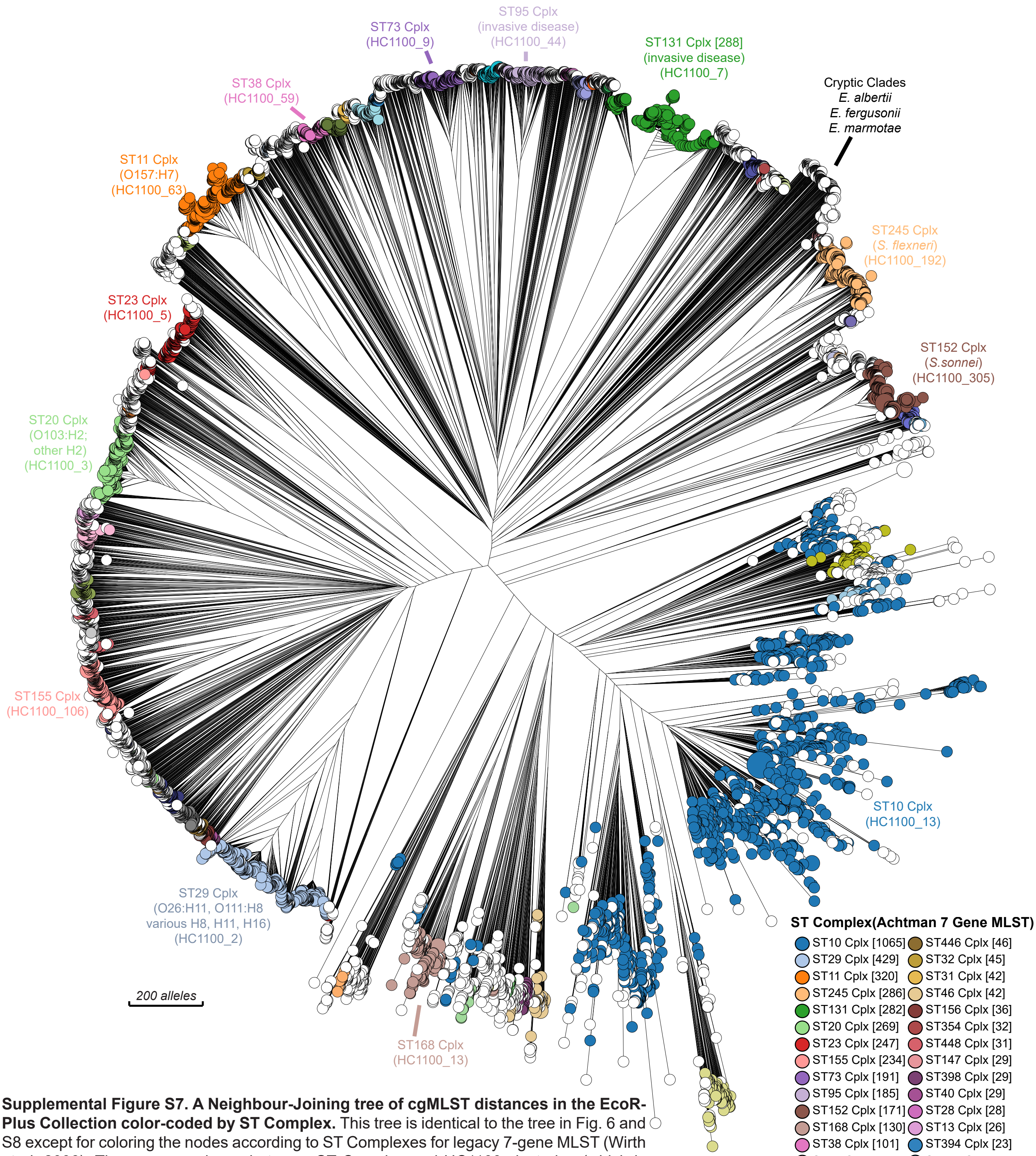


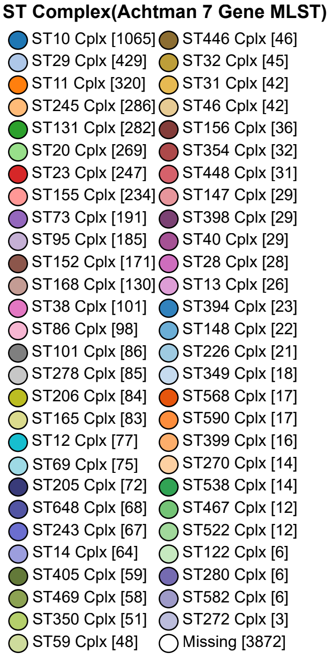


**Supplemental Figure S7. A Neighbour-Joining tree of cgMLST distances in the EcoR-Plus Collection color-coded by ST Complex.** This tree is identical to the tree in Fig. 6 and S8 except for coloring the nodes according to ST Complexes for legacy 7-gene MLST (Wirth et al. 2006). The correspondence between ST Complex and HC1100 clustering (which is based on much higher resolution cgMLST) is striking for the most common ST Complexes, with the exception of ST168 Complex (07:00) which is assigned to HC1100_13 by Hierarchical Clustering, and additional, rarer ST Complexes. The discrete nature of multiple ST Complexes according to cgMLST is noteworthy, and prominent examples of such discrete Complexes are indicated by text. Recent publications have provided interesting details on the ST131 Complex (usually erroneously referred to as ST131) (Johnson et al. 2016; Stoesser et al. 2016; Ben Zakour et al. 2016; Liu et al. 2018), the ST95 Complex (Achtman et al. 1983; Wirth et al. 2006; Gordon et al. 2017) and ST11 Complex/O157:H7 (Leopold et al. 2009; Dallman et al. 2015). Recent attention based on genomes in EnteroBase has been dedicated to ST1193 of ST14 Complex (Johnson et al. 2019). However, little attention has been directed at the other ST Complexes although they are a very common cause of disease in humans and animals according to the frequencies of their genomes in EnteroBase.

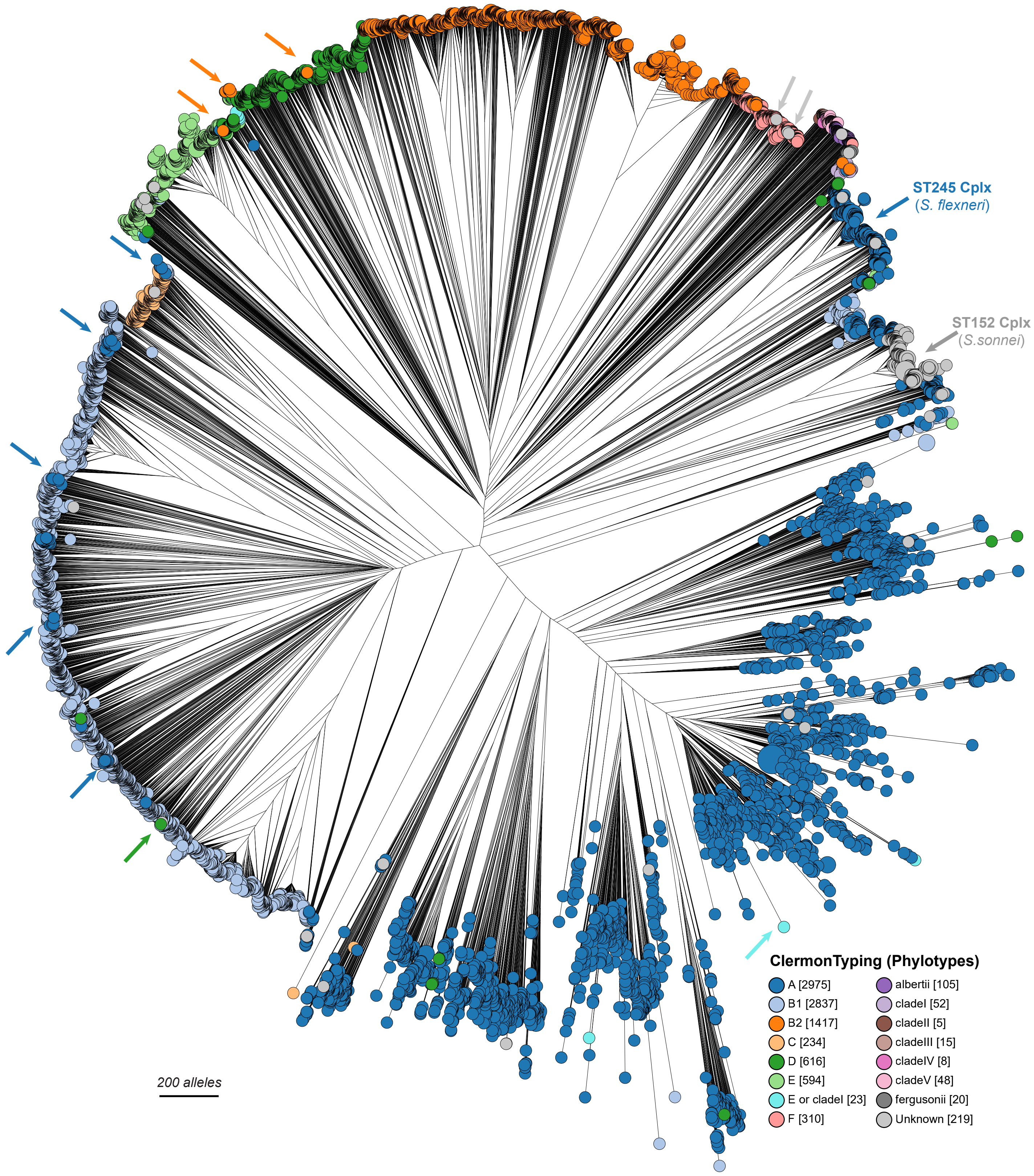


**Supplemental Figure S8. A Neighbour-Joining tree of cgMLST distances in the EcoRPlus Collection color-coded by Clermont types.** This tree is identical to the tree in Fig. 6 and Fig. S7 except that the nodes show the Clermont Types predicted by the program ClermonTyping (Beghain et al. 2018), which has been implemented within EnteroBase. Large parts of the tree are relatively homogeneous, indicating that Clermont Typing often correlates well with HC2000 clustering. However, arrows indicate multiple nodes which differ in Clermont Type from their close neighbors, illustrating that the presence/absence of genes from the accessory genome which is used for the Clermont scheme does not correlate completely with the phylogenetic relationships revealed by cgMLST. As a result, nodes assigned to Clermont Types A and B2 are found at multiple positions within the tree, far from most other strains of those Clermont types. In addition, two groups of Shigella are inaccurately labelled by Clermont Types. ST245 Complex largely corresponds to *Shigella flexneri* (Wirth et al. 2006), but is inappropriately assigned to Clermont Type A. Similarly, ST152 Complex largely corresponds to Shigella sonnei but is not recognized by Clermont Typing.

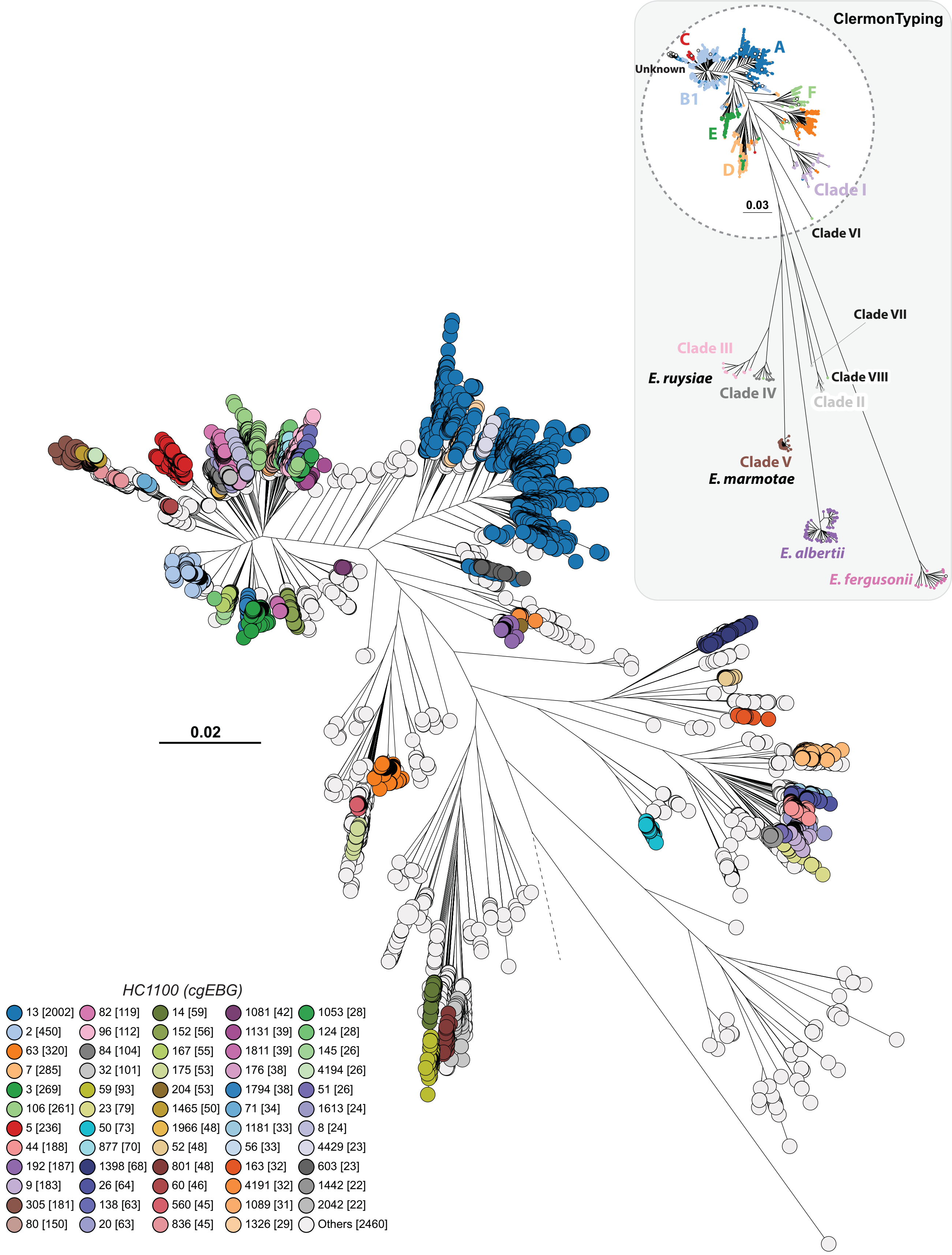


**Supplemental Figure S9. A Maximum-Likelihood (ML) tree of the EcoRPlus Collection.** 1,230,995 core SNPs were extracted from 9,479 core genomes after concatenating the sequences (2.33 Mbps) of their 2,513 core gene alignments with MAFFT (Katoh and Standley 2013). A maximum likelihood tree was calculated using FASTTREE 2 (Price et al. 2010). **Inset**) The ML tree of all genomes color-coded by ClermonTyping, including the Cryptic Clades I-VI, two novel cryptic clades (VII-VIII) and the *Escherichia* species *albertii, fergusonii*, and *marmotae*. Note that the three genomes of *E. marmotae* are on a deep branch within Clade V, in agreement with independent recent observations (van der Putten et al. 2019), which also assigned Clades III plus IV to the novel species *Escherichia ruysiae*. The white circle encloses genetically-related populations within *E. coli*, including Clade I, whereas the other Clades and species are on external branches within the gray rectangle. These topological relationships are similar to those described on smaller datasets (Luo et al. 2011; van der Putten et al. 2019). **Main figure**) Closeup of genomes on branches within the inner circle of the inset, color-coded by HC1100 HierCC cluster. This ML clustering of individual genomes is concordant with the clustering according to the neighbour-joining algorithm in Fig. 6, but provides much more accurate branch lengths. However, this tree also took several weeks to complete whereas Fig. 6 was complete in less than an hour.
